# MTALTCO1: a 259 amino-acid long mtDNA-encoded alternative protein that challenges conventional understandings of mitochondrial genomics

**DOI:** 10.1101/2024.09.22.614361

**Authors:** Francis Robitaille, Amy Campbell, Aziz Ben Hadj, Sarah Cornet, Ludovic Nadeau-Lachance, Thierry Niaison, Thierry Choquette, Liam Nyffeler, Xavier Roucou, Annie Angers, Sophie Breton

## Abstract

Mitochondrial alternative Open Reading Frames (ORFs) substantially broaden the functional scope traditionally attributed to mitochondrial DNA, encoding peptides and proteins that participate in diverse cellular processes. These newly identified ORFs are embedded within annotated sequences, both coding and non-coding, and reveal layers of overlapping genetic information. We here report the discovery of MTALTCO1, a 259 amino-acid protein, the longest mitochondrial alternative protein identified to date, encoded by an ORF located within the human cytochrome oxidase 1 gene, in the +3 reading frame. We confirm the expression and mitochondrial origin of MTALTCO1 through multiple independent lines of evidence, including a custom-designed antibody, mass spectrometry-derived peptides, sequence analysis, and inhibitors of mitochondrial expression. Despite encoding AGR codons as arginine, contrary to the prevailing view that these function as stop codons in the vertebrate mitochondrial genetic code, MTALTCO1 shows strong evidence of mitochondrial translation, challenging established models of mitochondrial codon usage and gene expression. Co-immunoprecipitations and pull-down assays delineate MTALTCO1’s interaction landscape across major cellular pathways. Lastly, we present the first in-depth analysis of conservation for a mitochondrial alternative ORF overlapping a reference protein-coding gene and discuss the results in light of MTALTCO1’s suggested role in protein scaffolding.

## INTRODUCTION

Human mitochondria rely on a proteome of over 1,100 proteins, with expression varying depending on cell types and physiological conditions (Morgenstern et al., 2021). Although most of these proteins are encoded by nuclear genes, translated in the cytoplasm and subsequently imported into mitochondria (Morgenstern et al., 2021; Palmfeldt & Bross, 2017), the mitogenome contributes to a small but essential portion of their resident proteome (Adams & Palmer, 2003; Butenko et al., 2024). The human mitogenome is a circular DNA molecule (mtDNA) of 16,569 base pairs commonly thought to encode 37 genes: 13 produce key subunits of the electron transport chain and ATP synthase, two code for ribosomal RNAs (12S and 16S rRNAs), and 22 encode transfer RNAs (tRNAs) that decipher the mitochondrial genetic code (Anderson et al., 1981; Rackham & Filipovska, 2022). This view was challenged in 2001 with the discovery of *humanin*, a small open reading frame (ORF) within the 16S rRNA gene encoding a 24 amino acid (AA) peptide (Hashimoto et al, 2001). Humanin revealed the first of a novel class of mitochondrial-derived peptides, termed MDPs, encoded by previously unrecognized ORFs nested in or overlapping annotated coding and non-coding regions of the mitogenome (Lee et al., 2013). Humanin has been characterized extensively and was shown to exert diverse biological effects, including but not restricted to neuroprotective and anti-apoptotic roles (Nishimoto et al., 2004; Charununtakorn et al., 2016; Yen et al., 2013; Lei & Rao., 2022; Boutari et al., 2022). By 2016, additional MDPs were identified in rRNAs, including MOTS-c and small humanin-like peptides (SHLP 1-6), ranging from 16 to 38 AA in length (Lee et al., 2016; Cobb et al., 2016). MOTS-c’s direct application can mimic exercise by regulating nuclear gene expression, enhancing glucose metabolism and metabolic homeostasis. Other SHLPs influence apoptosis, insulin sensitivity, and metabolic regulation, among other things (Kim et al., 2017). Evidence of purifying selection acting on Humanin and SHLP6 further supports their physiological relevance (Gruschus et al., 2023).

MDPs had until recently been primarily investigated in non-coding mitochondrial genes, where selection pressures offer an environment more conducive to the evolution of alternative protein function than coding regions (Gruschus et al., 2023). The discovery of SHMOOSE in 2023 (Miller et al., 2023), a 58 AA protein whose coding sequence overlaps tRNA Serine and ND5, marked the onset of a series of increasingly larger mitochondrial-derived proteins found within coding regions. SHMOOSE was shown to influence mitochondrial gene expression and regulate oxygen consumption (Miller et al., 2023). The same year, MTALTND4, a 99 AA protein –larger than some canonical mitochondrial proteins like ATP8 and ND4L– was identified as an alternative ORF nested in the ND4 gene. MTALTND4 induced a quiescent state-like metabolic depression when exogenously administered to HeLa and HEK-293T cells (Kienzle et al., 2023). In 2024, CYTB-187AA, a 187 AA protein whose coding sequence is embedded within the *cytochrome b* gene, was shown to promote pluripotency during the primed-to-naive transition in embryonic stem cells (Hu et al., 2024).

In light of the growing recognition that mitochondria are active organelles that perform functions extending far beyond energy production (Monzel et al., 2023; Picard & Shirihai, 2022; Spinelli & Haigis, 2018; Picard & Sandi, 2021), alternative proteins encoded by the mitogenome warrant particular attention for their potential to expand the organelle’s functional repertoire and challenge the long-standing view of mitochondrial genomes as functionally frozen, at least in Metazoans (Shtolz & Mishmar, 2023). Their expression, either fully or partially within the mitochondrion itself, imparts a level of organelle-specific agency not shared by nuclear-encoded proteins, and may be especially relevant in the context of mitonuclear and inter-organelle crosstalk, where targeted and localized regulation and signaling dynamics are critical (Picard & Shirihai, 2022). Mitochondrial-derived alternative proteins’ emerging relevance to human health, spanning from biomarker discovery (Rochette et al., 2022; Cai et al., 2022) to disease aetiology (Miller et al., 2022; Kim et al., 2021) and therapeutic target development (Dabravolski et al., 2021; Arneson et al., 2022), further underscores their importance.

Motivated by their emerging biomedical relevance and potential to expand the functional scope of the mitogenome, we confirm the existence and mitochondrial origin of MTALTCO1, a 259 AA alternative mitochondrial protein whose coding sequence is nested within the CO1 gene.

## MATERIAL AND METHODS

### Custom rabbit anti-MTALTCO1 antibody generation

Kienzle et al. (2023) interrogated the CPTAC_Breast_Cancer database, a large repository of more than 32 million mass spectrometry MS/MS scans. Using the PepQuery tool, they retrieved peptide-spectrum-matches (PSM) scores of peptides drawn from an in-silico digestion of the theoretical mitochondrial alternative proteome generated with the mitochondrial genetic code using AGG and AGA (AGR) as arginine. A MTALTCO1-derived peptide (ELLPPWSLR) was identified and a custom rabbit anti-MTALTCO1 (antigenic peptide PNAPLRLIRPNHSSPTSPISPSPSCWHHYTTN) was ordered from MediMabs (Montreal, QC, Canada). The signal obtained was of expected weight, but did not disappear in Rho0 cells lysates obtained from Abcam (ab154479), casting doubt on its possible mitochondrial origin (Kienzle et al. 2023). In the present study, osteosarcoma 143B-ρ0 cells (Kerafast [ESA106]) were used, together with chloramphenicol and actinonin treatment in other cell types, to show that MTALTCO1 derives from mitochondrial DNA (see below).

### Cell culture and treatments

HEK-293T and HeLa cells were cultured in Dulbecco’s modified eagle medium (DMEM) supplemented with 10% of fetal bovine serum (FBS), antibiotics (1% streptomycin and penicillin) in 10 cm petri dishes maintained at 37°C with 5% CO_2_. Osteosarcoma 143B-ρ0 cells were grown in the same medium but supplemented with 1mM pyruvate and 5 µg/µL uridine. Actinonin treatment was realized on HeLa cells split into 2 groups, one serving as control and left untouched for 48h, and the other incubated with actinonin (150 µM) for 48h. Chloramphenicol treatment was realized on HeLa cells as in Kienzle et al. (2023), i.e. by adding 2 mL of chloramphenicol (50 µg/mL in 95% ethanol) in a 10-cm petri dish of 80% cell confluency. Cells were treated for 48h.

### Protein extraction and Western Blot

The full western blot protocol can be found in supplementary material (Supporting Text 1). Briefly, cells were washed, harvested and sonicated, and proteins were extracted as in Kienzle et al. (2023). Total protein concentration was determined using the Bradford assay, and aliquots containing 100 µg of proteins were stored at -80°C for later use. Various tissue lysates (kidney, heart, testis, brain, liver, skeletal muscle, spleen) were also obtained from Novus Biologicals (NB820-59231; NB820-59217; NB820-59266; NB820-59177; NB820-59232; NB820-59253; NB820-59259). Protein extracts were separated by SDS-PAGE before being transferred to a nitrocellulose membrane for blotting with primary and secondary antibodies. The primary antibodies used were: “rabbit anti-MTALCO1” (1:1000) (MediMabs), “mouse anti-ATP5” (1:2000) (Abcam [ab14748]), “mouse anti-CO1” (1:1000) (LifeSpan BioSciences, Inc. [LS-C172110]), “rabbit anti-ND4” (1:1000) (Boster Biological Technology Co., Ltd. [A04180-1]), and “mouse anti-actin” (1:2000) (Abcam [ab1801]). The secondary antibodies used were goat anti-rabbit IgG (1:2000) or goat anti-mouse IgG (1:2000) coupled to horseradish peroxidase (HRP). The specificity of our anti-MTALTCO1 has also been verified by adding the synthetic antigenic peptide to the primary antibody solution at a 10 X concentration before incubation. This step was performed to competitively chelate every antigenic site of the primary antibody, confirming the attenuation of the specific MTALTCO1 band. Revelation was achieved by chemiluminescence using West-Pico SuperSignal (Thermo Fisher Scientific).

### Immunofluorescence

The full immunofluorescence protocol can be found in supplementary material (Supporting Text 1). Briefly, cells were cultured on coverslips coated with poly-D-Lysine and then fixed and permeabilized as in Kienzle et al. (2023). After a 30-minutes blocking step at room temperature, a 2-hours incubation was performed with 1:50 rabbit anti-MTALTCO1 and 1:250 mouse anti-ATP5 primary antibodies in PBS (1% NGS) (Kienzle et al. 2023). After washes, the cells were incubated with secondary IgG antibodies (respectively Alexa Fluor TM 488 anti-rabbit [1:500] and Alexa Fluor TM 488 anti-mouse [1:500]) in PBS (1% NGS) for 1 hour at room temperature. Finally, the cells were washed with PBS and mounted in DAPI mounting medium, then visualized using a Zeiss-LSM800 confocal microscope.

### Pull-down, Co-Immunoprecipitation and Mass Spectrometry

Full protocols of cell lysate extractions, pulldown assays and co-immunoprecipitation assays can be found in supplementary material (Supporting Text 1). In short, GST pull-down assays and co-immunoprecipitations were performed as in Kienzle et al. (2023) with slight modifications. For the pull-down experiments, GST alone was used as a control and the fusion protein GST-MTALTCO1 was used to isolate possible interacting partners. These proteins were immobilized on Pierce glutathione magnetic agarose beads (Thermo Scientific). For the co-immunoprecipitation assays, Protein A Mag Sepharose magnetic beads (Cytiva) were added to the custom rabbit anti-MTALTCO1 antibody and as a control, the rabbit pre-immune serum was used. The antibodies were crosslinked to the magnetic beads as described by the manufacturer. Following incubation with cell-lysates, magnetic beads were washed and resuspended in Tris-HCl and submitted to the proteomics core facility of the Institute for Research in Immunology and Cancerology (IRIC, University of Montreal) for mass spectrometry analysis.

The resulting data were analyzed with Scaffold 5.0, using a 99% protein threshold and 95% peptide threshold. Total spectrum counts and exclusive unique peptide count were determined for each protein detected in both experimental and control samples. Proteins identified in both experiments were filtered and compiled (Supplementary Material: Supporting Data 1). Filtered proteins were analyzed by the software STRING (string-db.org), to determine enriched pathways MTALTCO1 may be involved in (Szklarczyk et al., 2025).

### Pseudogene screening

CO1 pseudogene nucleotide sequences, derived from GenBank’s web server on account of nucleotide similarity with our antigenic peptide sequence, were aligned in MEGA using the muscle algorithm (Stecher et al., 2020), and translated using the standard genetic code.

### Sequence analysis: structure and disorder

Structure predictions were made on the AlphaFold’s web-server (Abramson et al., 2024). Intrinsic disorder predictions for known Mitochondrial alternative ORFs (MtaltORFs) of length greater than 31 AA, the 1136 proteins listed in MitoCarta (Rath et al., 2021), and 100 simulated degenerate sequences (see below) were made on the RIDAO web-server (Dayhoff & Uversky, 2022), using the VSLB2 score as per Nesterov et al., (2024).

### Conservation of MTALTCO1

PhyloP conservation scores for 30 primates were obtained from the UCSC Genome Browser. In our first analysis, these scores were smoothed using a moving average filter, with a sliding window of 25 nucleotides. We then separated nucleotides in a codon-based logic, with first, second and third nucleotides in the CO1 reading frame being grouped together and compared for their PhyloP scores.

To compare positional information conservation at the amino acid level between CO1 and MTALTCO1, we retrieved CO1 nucleotide sequences from 27 primates, aligned them, and constructed a phylogenetic tree using the Tamura-Nei substitution model under a Maximum Likelihood framework in MEGA (Stecher et al., 2020). The resulting tree was used to define a linear ordering of primates from 0 to 26, where 0 refers to *Homo sapiens* (full list can be found in supplementary file, supporting text 3). This ordering minimizes neighbor distances, thereby providing a conservative estimate of neighbor divergence. Each aligned sequence was then translated into its corresponding protein sequence, using the +1 reading frame for CO1 and the +3 frame for MTALTCO1. Pairwise Hamming distances were computed between the amino acid sequences and graphed using a color scale to reflect sequence divergence.

### Comparing MTALTCO1 to random degenerate sequences

To generate random nucleotide sequences encoding reference proteins while conforming to specific sequence features observed across primates, we first gathered statistics of various sequence features (stop codon content in alternative frames, CpG and GC content). We then instantiated a randomly generated nucleotide sequence encoding the CO1 gene, constrained only by the exclusion of AGR codons in the reference frame. A random walk introduced single-codon mutations at each step, after which sequence features were evaluated and compared to the defined target distributions. Mutations that moved the sequence closer to the target ranges were retained. This process was iterated until all features fell within their respective target intervals, at which point the sequence was accepted. The resulting ensemble of such sequences is referred to as the Degenerate Sequence Space (DSS).

One thousand such sequences were generated for *Homo sapiens*, and the lengths of ORFs spanning the region between the start and end positions of MTALTCO1 were collected in the +3 reading frame. Then, 20 DSS sequences were generated per primate species (n = 27) and compiled in a single FASTA file containing real and DSS sequences. For each sequence in the +3 reading frame, both real and simulated, protein molecular features (see localCIDER for additional information on molecular features, Ginell & Holehouse, 2020) and amino acid usage were computed, normalized across all sequences, and subjected to principal component analysis (PCA). To characterize the distribution of generated sequences in the reduced feature space, a kernel density estimator (KDE) was applied to the PCA scatterplot. Density was visualized via topographic contour lines and color-coded according to local point density. Each degenerate sequence was assigned a density value from the KDE output and ranked from least to most dense in this projected space. Real protein sequences were mapped onto this density landscape and ranked according to their positions in the DSS-ranking. For instance, a sequence ranked beyond the 99.9% threshold is assumed more unlikely to happen at random than 99.9% of the generated sequences in this reduced space. This non-parametric approach was chosen to avoid assumptions about the underlying density distribution, particularly given the increased granularity and radial asymmetry away from the distribution’s center.

### Preservation of MTALTCO1 in human lineages

We interrogated every mitochondrial variant in the MitoMap (Rath et al., 2021) database to assess whether MTALTCO1 is preserved in major mitochondrial lineages. Each variant was mapped onto the revised Cambridge Reference Sequence NC_012920. Variants were classified as destructive if they either disrupted the annotated start codon or introduced premature stop codons. All destructive variants were compiled for major mitochondrial haplogroups.

We then adapted a publicly available constraint analysis from (Lake et al., 2024), which constructs a mutational model to assess constraint (the removal of deleterious mutations at the population level) by comparing observed-to-expected ratios for synonymous, missense, and stop-gain mutations. After running their analysis, we extended it to include a constraint assessment of alternative proteins overlapping canonical protein-coding genes. In this extension, we focused on mutations that are synonymous in the reference ORFs, allowing us to derive a constraint metric specific to those overlapping ORFs. These analyses were run on the *gnomAD* dataset, and consider heteroplasmy levels of mitochondrial variants.

## RESULTS & DISCUSSION

### Endogenous detection of MTALTCO1 in immortalized cell-lines and tissue lysates

MTALTCO1 (Mitochondrial Alternative CO1), in keeping with the nomenclature introduced by Kienzle et al. (2023), was confirmed as a 259 AA protein whose 777 base pair long coding sequence nests in the mitochondrial cytochrome oxidase 1 gene in the sense +3 reading frame, i.e. shifted by two nucleotides downstream (Figure 1A). The signal detected for the protein (∼ 31 kDa) with our custom antibody in HEK-293T and HeLa cells corresponds to the expected molecular weight (Figure 1B). MTALTCO1 exhibits a tissue-specific expression pattern, showing the highest expression level in the kidneys, testis, and spleen (Figure 1E). In contrast, heart and skeletal muscle tissues show only faint expression, whereas the liver and brain display little to no detectable expression. Confocal microscopy data suggests that MTALTCO1 might localize in the cytoplasm (Figure 1G).

**Figure 1.**
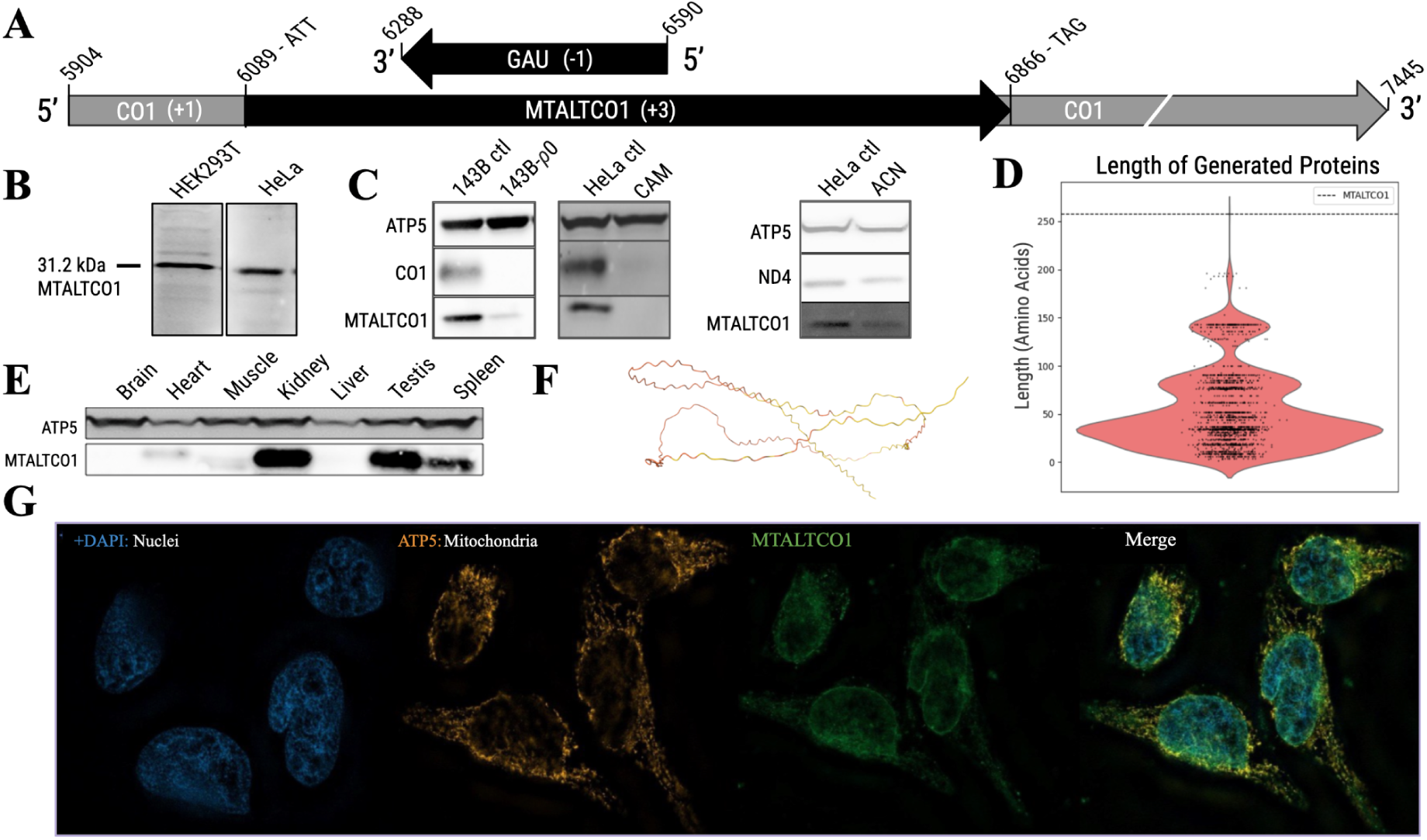
MTALTCO1 is a 259 amino acid protein encoded in the +3 reading frame of the CO1 gene. (A) MTALTCO1 is initiated by ATT and terminated by TAG, translated from the mitochondrial genetic code to produce a 259 AA long protein. CO1 is located in the +1 frame (reference frame), MTALTCO1 in the +3 reading frame. (B) MTALTCO1 detected by Western Blot in HEK293T cells as well as in HeLa cells. (C) Proof of mitochondrial origin: ATP5 is the nuclear control, CO1 and ND4 are mitochondrial controls. 143-B ρ0 cells, chloramphenicol (CAM) and actinonin (ACN) are inhibitors of mitochondrial expression. (D) Density plot of the lengths of 1000 DSS-derived proteins for *Homo sapiens*. The dashed line marks the length of MTALTCO1. (E) MTALTCO1 and nuclear-encoded mitochondrial ATP5 detection by Western Blot in various tissue lysates. (F) ALphaFold2 prediction of structure. Orange: very low confidence, yellow: low confidence. (G) MTALTCO1 and nuclear-encoded mitochondrial ATP5 detection by immunofluorescence in HeLa cells stained with DAPI.

LC-MS/MS analysis of proteins immunoprecipitated by the anti-MTALTCO1 antibody identified multiple peptides matching the MTALTCO1 sequence, resulting in a protein identification with 100% confidence. In total, 30 total spectra were detected (with 20 exclusive unique peptides), covering 54% of the MTALTCO1 amino acid sequence (Figure 2A). We interrogated all nuclear CO1 pseudogenes to see if they could explain some of our peptide matches and found that while there were some sequence matches, none had upstream cytoplasmic initiation codons (Supplementary Material: Supporting Text 2 and Figure S1). Additionally, four unique peptides who had no pseudogene equivalent in sequence were identified, three with 100% confidence and one with 99% confidence, further making pseudogenes an unlikely confounding factor in the identification of MTALTCO1. In support to this is the observation that none of the numerous CO1-derived NUMTs code for a complete MTALTCO1 nor provide an antigenic sequence that is not immediately flanked by stop codons (Supplementary Material: Supporting Text 2 and Figure S1).

**Figure 2.**
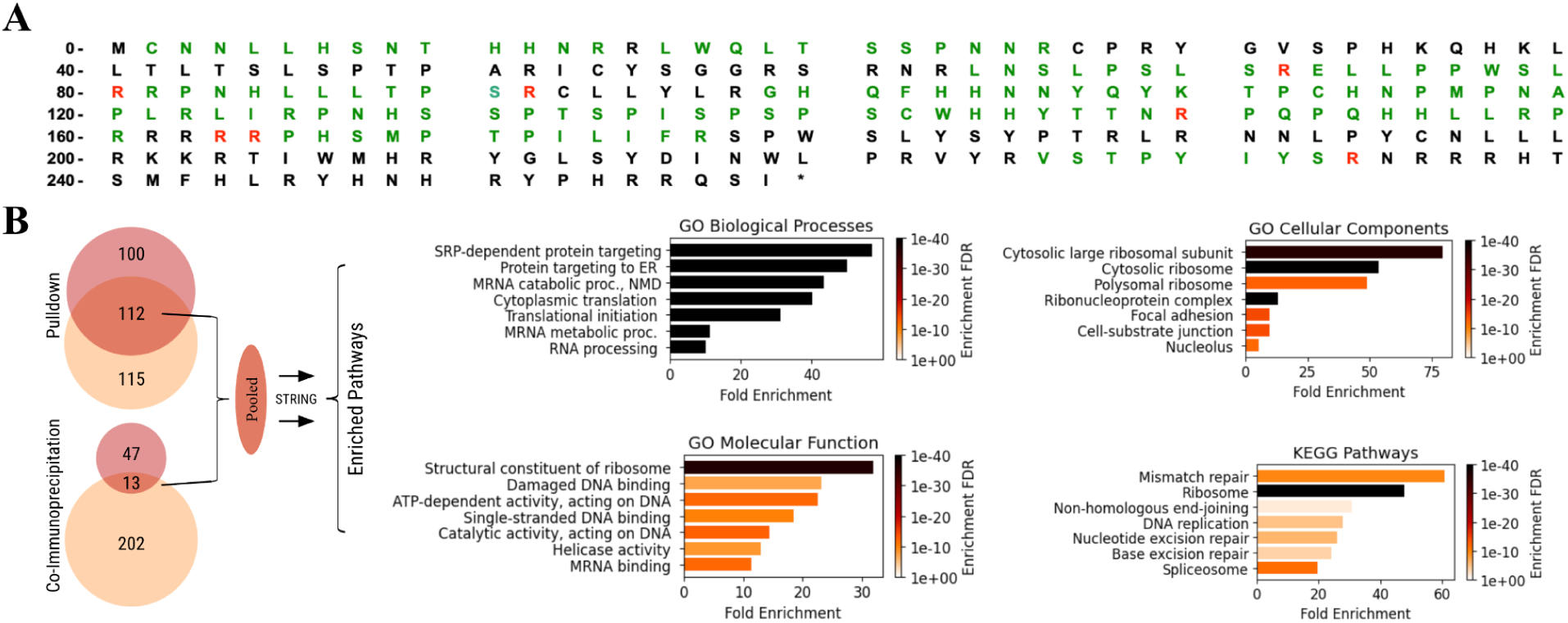
MTALTCO1 sequence coverage in MS/MS spectra and enriched pathways. (A) The amino acid sequence coverage for MTALTCO1 is 54%, for 30 total spectra, 25 exclusive unique spectra and 20 exclusive unique peptides. The amino acids in green and red are present in at least one spectrum, with the latter being arginine residues coded by AGA/AGG. (B) Count of enriched proteins per interactor studies shown on the left, with top 10 enriched pathways per pathway annotation pipeline shown on the right.

Of particular interest is the observation that the *Gau* open reading frame (ORF), originally described by Faure et al. (2011), is entirely embedded within the *MTALTCO1* sequence, which itself is nested within the canonical *CO1* gene. *Gau* is translated in the –1 reading frame relative to *CO1*, such that their codons share the same third (wobble) nucleotide positions. As a result, *Gau* displays predominantly reciprocal synonymy with the *CO1* frame, effectively mirroring the latter’s high degree of amino acid conservation. Notably, this arrangement minimizes competition with *MTALTCO1*, with only limited entanglement of codon constraints at the third nucleotide position. This configuration results in a unique genomic region encoding three fully overlapping ORFs, a remarkable and, to our knowledge, unprecedented level of information density within a single sequence. The functional and evolutionary implications of this intricate genomic entanglement warrant further investigation.

### MTALTCO1 is translated in the mitochondria

Unlike most known MDPs, which typically retain cytoplasmic initiation codons, MTALTCO1 lacks any canonical cytoplasmic start codons, reducing the likelihood of cytoplasmic translation. Even under the hypothetical scenario of alternative initiation codons instigating cytoplasmic translation of an exported transcript, MTALTCO1 contains a TGA encoded tryptophan (cytoplasmic stop codon and assigned to tryptophan in the mitochondrial code), which would break the ORF and produce a truncated protein of at most 191 amino acids. However, such bands were not detected in our western blot analysis. Furthermore, MS-peptides KTPCHNP<u>M</u>PNAPLR and RRPHS<u>M</u>PTPILIFR contain methionines encoded by ATA codons in the MTALTCO1 nucleotide sequence (Figure 2A), consistent with the well-established reassignment of ATA from isoleucine to methionine in vertebrate mitochondria. Lastly, we assessed loss-of-signal following inhibition of mitochondrial expression. In 143-B *ρ*0 cells, which lack endogenous mtDNA, a loss-of-signal was observed. Cells treated with chloramphenicol, which inhibits protein synthesis in mitochondria, similarly showed a loss of signal. Cells treated with actinonin, which inhibits mitochondrial transcription, showed a reduction in signal (Figure 1C).

### The translation of MTALTCO1 challenges current models of mitochondrial DNA expression

Multiple independent lines of evidence support the existence and mitochondrial origin of MTALTCO1. Confidently establishing both is particularly important, given their potential to challenge the prevailing view that AGA and AGG (AGR) codons serve exclusively as stop codons in the mitochondrial genetic code due to the absence of corresponding tRNAs (Jukes & Osawa, 1990). AGR codons seem to be decoded as arginine in MTALTCO1 (labelled in red in Figure 2A), and along with others (Faure et al., 2011; Seligmann 2011 and 2012; Kienzle et al., 2023), we suggest that these codons may be functionally reassigned to code for arginine in mitochondrial alternative reading frames. In further support of this reinterpretation, the unique MS-peptide <u>RR</u>PHS<u>M</u>PTPILIFR is coded by an ATA codon decoded as methionine (underlined), consistent with the mitochondrial genetic code, and also by two AGR codons decoded as arginine residues (also underlined). This peptide could not arise under either the universal genetic code nor the reference mitochondrial code unless AGR codons are reassigned, reinforcing the case for codon reassignment in alternative frames. Given the enrichment of AGR codons in the +3 alternative frame, the existence of an overlapping genetic code is non-trivial, as it may reveal underlying regulatory logics of the alternative proteome (Seligmann, 2012) and inform functional hypotheses. A comprehensive discussion on the reassignment of AGR codons is provided in Supplementary Material (Supporting Text 2).

Recent models of mitochondrial gene expression models tend to overlook the various complex challenges posed unto them by mtaltORFs and might therefore fail to fully capture their unique dynamics (Rackham & Filipovska, 2022). We propose that the identification and reliable detection of these mtaltORFs, especially those challenging established principles paradigms, will provide valuable insights to improve and refine current models.

### MTALTCO1’s unusual length suggests functional relevance

The degeneracy of the genetic code has been leveraged to identify functional overlapping ORFs, by comparing their observed lengths to the expected lengths of ORFs generated from random sequences constrained by the amino acid sequence of the annotated host ORF. Such an approach, previously applied in the identification of overlapping ORFs in viral genomes (Schlub et al., 2018), is particularly effective for detecting functional ORF overlaps exceeding 50 nucleotides, a relevant parallel given that both viruses and mitochondria overprint in response to stringent genomic packing constraints (Pavesi, 2021; Brandes & Linial, 2016). MTALTCO1 stands out as an outlier in the length distribution of generated degenerate ORFs in the +3 frame of CO1 (Figure 1D), suggesting a depletion of canonical stop codons in this reading frame. This is notable because mitochondrial alternative frames are not expected to be neutrally populated with stop codons; rather, according to the ambush hypothesis, they should be actively enriched to prevent spurious translation events (Seligmann, 2010).

We then monitored whether the full-length ORF of MTALTCO1 is preserved at the population level and found that it is preserved in more than 99.28% of all sequences in the MitoMap database (Rath et al., 2021), comparable to the 99.4% preservation observed for Humanin (Logan, 2017). This occurs despite MTALTCO1’s much longer length, which provides more mutational opportunities for premature stop codons to arise by chance. Variants assigned to the N haplogroup preserved MTALTCO1 in at least 99.9% of sequences, those in the M haplogroup in over 99.7%, and those in the L haplogroup in more than 94.8%. Variants that disrupted the MTALTCO1 start codon were absent from at least 99.99% of all sequences. A nonsense variant (Q154*) that pseudogenized MTALTCO1 and was present in 5.19% of sequences within the L haplogroup, accounted almost entirely for the reduced preservation observed in this lineage.

To verify whether this preservation is simply contingent on scarce mutational opportunities to break MTALTCO1’s ORF, we adapted a public constraint analysis (Lake et al., 2024) that monitors the removal of mitochondrial mutations at the population level. Our analysis indicates strong constraint against stop-gain mutations in MTALTCO1. However, this signal of constraint cannot be disentangled from constraint on missense mutations in the reference reading frame, as it disappears when the analysis is restricted to mutations that are synonymous in CO1 (Figure 3E). Constraints on stop gain variants may thus reflect constraint entanglement across overlapping frames, highlighting the need for further work to investigate the implications of such inter-frame constraints.

**Figure 3.**
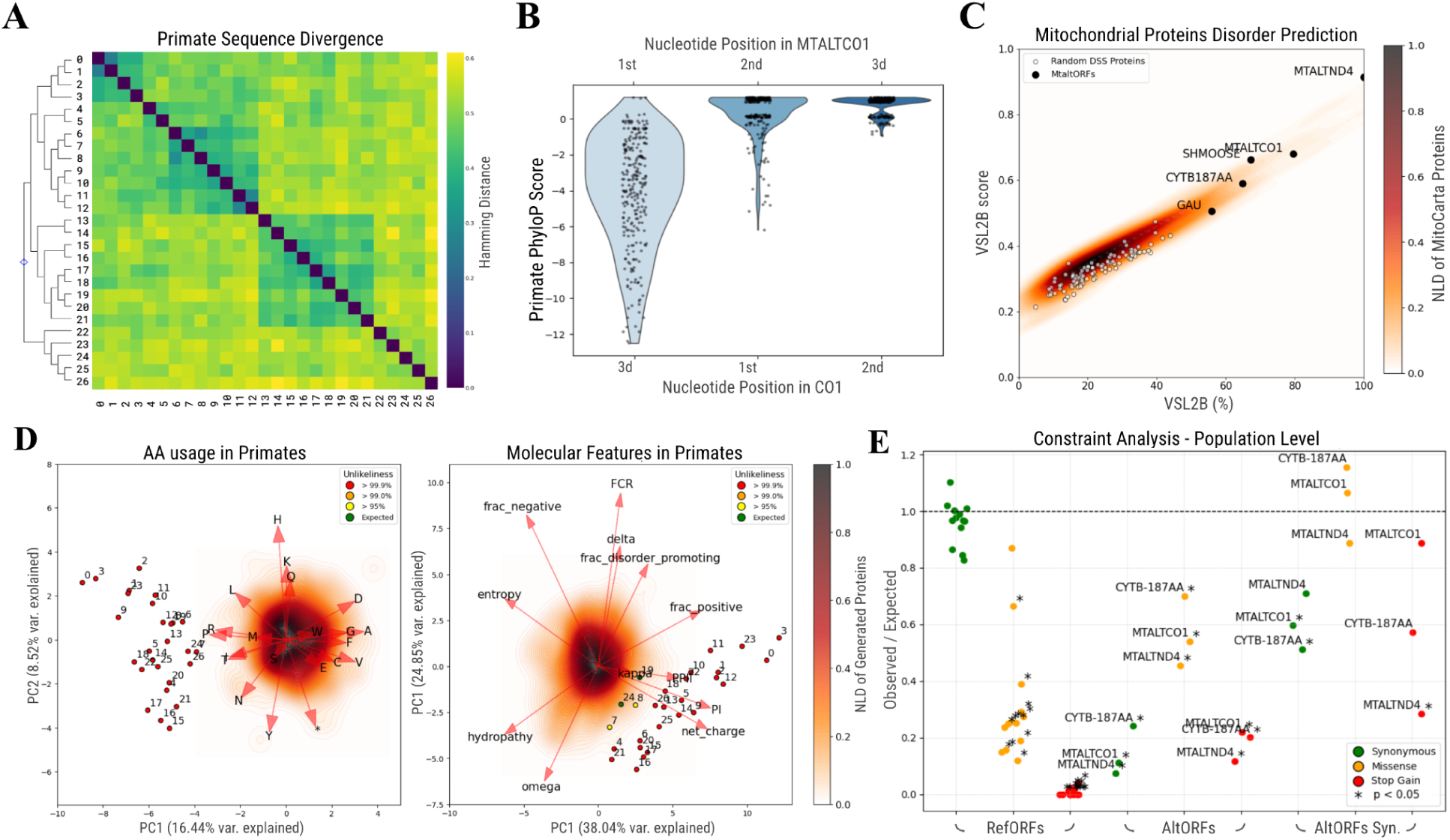
Various bioinformatic metrics for MTALTCO1. (A) Pairwise Hamming distances between primate MTALTCO1 sequences, with species grouped and ordered from 0 (*Homo sapiens*) to 26. Full species list provided in Supplementary File 2. (B) PhyloP nucleotide conservation scores, ordered according to the RF of either CO1 or MTALTCO1. (C) Disorder predictions for the entire mitoproteome using the VSL2Bdisorder prediction algorithm. Normalized Log Density (NLD) color-codes the density distribution of the human mitoproteome. Disorder scores for MDPs over 31 AA in lengths are overlaid and labelled, along with generated CO1-DSS proteins. (D) Distribution of amino acid usage and molecular features in reduced feature space for generated (topographic contour lines - color-coded by the Normalized Log Density (NLD)) and real protein sequences (scatter - ranked and color-coded by their density percentile). Numbers correspond to species in panel A. (E) Constraint analysis, at the *Homo Sapiens* populational level, for three types of mutations. A significant result (*) means there is constraint. See methods for details.

The unusual length of MTALTCO1, substantially exceeding that expected from neutrally evolving overlapping ORFs, suggests functional relevance. To explore this possibility, we next examined its interaction partners, predicted structural features, and evolutionary conservation, using these complementary lines of evidence to guide functional inference.

### MTALTCO1 interacts prominently with ribonucleoprotein complexes and nucleic acids

To identify putative binding partners of MTALTCO1, we analyzed proteins co-immunoprecipitating with MTALTCO1 or selected by affinity binding to a GST-MTALTCO1 fusion protein. Among the identified candidates, 60 proteins passed the selection criteria in the co-immunoprecipitation experiment, while 227 proteins met the thresholds in the pull-down assay, with 10 being identified in both experiments (Figure 2B; Supplementary Material: Supporting Text 1 and Tables S1-3). The 277 proteins identified in the co-immunoprecipitation and pull-down experiments were submitted to functional enrichment analysis using the Gene Ontology (GO) and Kyoto Encyclopedia of Genes and Genomes (KEGG) databases. This analysis revealed significant enrichment of 177 GO terms related to Biological Processes, 35 terms associated with Molecular Functions, and 86 terms related to Cellular Components. In parallel, 12 KEGG pathways were found to be significantly enriched. The top 10 enriched GO and KEGG terms, ranked by Fold Enrichment, are presented in Figure 2B. Some of the proteins identified are nuclear, which is possible given the experimental context; however, MTALTCO1 is not observed in the nucleus in imaging data, and one would not typically expect it to interact with nuclear spliceosomes or other nuclear proteins. These interactions can nonetheless occur under the conditions used in co-immunoprecipitation and pulldown experiments.

Three major themes emerge from the enrichment analyses. First, MTALTCO1 is associated with cytoplasmic ribonucleoprotein (RNP) complexes, particularly structural components of the ribosome and the spliceosome. While cell lysis during interactome experiments can artifactually bring molecular components into proximity, the likely cytoplasmic localization of MTALTCO1 suggests that these associations may reflect physiologically relevant interactions. Second, enriched proteins point to broad interactions with nucleic acids, consistent with involvement in pathways related to RNA processing and splicing. Third, several enriched pathways are involved in genome integrity and maintenance, such as mismatch repair, non-homologous end joining, and base excision repair (Figure 2).

The elevated arginine content of MTALTCO1 (roughly 15%) concordantly suggests a high potential for nucleic acid binding. Arginine residues are known for their versatile “stickiness”, simultaneously engaging simultaneously in salt-bridges, hydrogen bonds, cation-π bonds, and hydrophobic interactions (Gupta & Uversky, 2023). They are notorious for their prominent role in mediating interactions with both proteins and nucleic acids, particularly with RNAs (Bayer et al., 2005). Whether our interactomics data reflect direct protein–protein interactions with RNP complex components, or are indirectly mediated via RNA, warrants further investigation.

### MTALTCO1 is an intrinsically disordered protein with conserved molecular features

Various evidence suggests that positional information in MTALTCO1 is poorly conserved. The considerable variability in MTALTCO1 sequences, both in relation to other primate species, and across neighboring species from proximal to distal order, contrasted with the higher conservation observed in the equivalent CO1 sequences (Figure 3A-B). Similarly, decomposed PhyloP scores across primates revealed a canonical conservation signature in the CO1 reading frame: the first and especially the second codon positions showed elevated conservation (Figure 3B). The third codon position, corresponding to a consequential position in the overlapping MTALTCO1 frame, exhibited the lowest average conservation. Our constraint analysis echoes these findings, suggesting no constraint on missense mutations in MTALTCO1 when controlling for shared constraint with the CO1 frame (Figure 3E).

Lack of positional conservation does not necessarily imply absence of functional potential, particularly in intrinsically disordered proteins (IDPs), where functional conservation often resides in molecular features rather than precise amino acid sequence (Zarin et al., 2021). MTALTCO1, in keeping with other MDPs larger than 31 AA, appears to exhibit greater intrinsic disorder than other mitochondrial proteins, with a VSLB2 score of 0.7, and a sequence disorder percentage of roughly 80% (Figure 1C). Perhaps counterintuitively, random sequences tend to exhibit less intrinsic disorder than natural proteins (Yu et al., 2016). To situate the disorder propensity of MTALTCO1, we compared it to a set of degenerate sequences derived from *Homo sapiens’* CO1 gene, and found it displayed a markedly higher level of intrinsic disorder than these controls (Figure 3C), reinforcing the idea that its disorder is non-random and potentially functional.

Furthermore, the functional relevance of MTALTCO1’s intrinsic disorder is supported by evolutionary evidence: while its positional conservation is rather poor, molecular features show evidence of selective constraint. When assessed against simulated degenerate primate sequences, MTALTCO1 and its primate orthologs stood out as clear outliers with respect to molecular features characteristic of IDPs (Ginell & Holehouse, 2020). Specifically, 25 out of 29 orthologs were more unlikely than 99.9% of simulated sequences, 2 exceeded the 95% threshold, and 2 sequences were within the expected range (i.e., within the central 95% of generated points; Figure 3D). Certain molecular features were consistently overrepresented across MTALTCO1 orthologs, including lower sequence entropy (low complexity), reduced negative fraction, and an elevated net charge. In line with findings from overlapping nuclear altORFs, we also observed a consistently higher isoelectric point (Vasu et al., 2022). Despite applying conservative tree-based grouping to minimize neighbor phylogenetic distance, the average pairwise dissimilarity between neighboring primate sequences remained high (mean = 42%; Figure 3A), suggesting that while these sequences have the mutational capacity to decay rapidly into the expected distribution, they instead retain certain unlikely molecular features.

### MTALTCO1 may contribute to subcellular organization as a scaffold protein

Structural disorder is a common feature of scaffold proteins, as it facilitates flexible and adaptable interactions with multiple binding partners (Buday & Tompa, 2010; Cortese et al., 2008). Scaffold proteins spatially and temporally coordinate sets of molecules by tethering them together (Good et al., 2011). More than passive linkers, scaffolds can actively modulate cellular functions, for example by tuning metabolic flux through scaffold titration or by modulating signaling pathways via controlled amplification (Good et al., 2011). When associated with mRNAs, scaffolds can act as central hubs for cellular signaling, integrating translation with broader regulatory processes (Smith & Costa, 2024). They can also contribute to the assembly and stabilization of enzymatic, signalization and ribonucleoprotein complexes (Cortese et al., 2008; Buday & Tompa, 2010).

Several features suggest that MTALTCO1 may act as a disordered scaffold protein capable of binding RNA, RNA-binding proteins, and/or ribosomal components. Central to this hypothesis is the interactor profile of MTALTCO1, which is predominantly associated with structural components and other constituents of the ribosome and translation machinery. As previously noted, the high overall arginine content in MTALTCO1 and its orthologs supports potential interactions with RNA and proteins. This enrichment, along with selective constraints favoring a high net positive charge and depletion of negatively charged residues, is consistent with affinity for nucleic acids. These features, conserved across species, suggest that such interactions may have evolutionary significance rather than being a species-specific or contingent property. Beyond the elevated arginine content itself, particular attention is warranted for its sequential arrangement in *Homo sapiens*, most notably a centrally located penta-arginine motif. Short arginine motifs are known to facilitate macromolecular assembly (Kyne & Crowley, 2017); for example, a tri-arginine motif mediates RNA binding on model protocell membranes (Kamat et al., 2015), while tetra-arginine motifs were sufficient to drive ‘promiscuous’ protein stickiness in the *E. coli* cytoplasm (Kyne & Crowley, 2017). Some RNA-binding proteins are known to interact with RNA in a sequence-independent manner, thereby aligning with the limited positional conservation observed across various scales, suggesting that the high sequence variability may reflect neutral drift constrained only by the preservation of molecular features compatible with RNP or RNA binding (Lukong et al., 2008).

Further work is required to validate roles in protein scaffolding, characterize the specificity and valency of these interactions, and elucidate their functional consequences within the cellular context.

### Interpreting the dynamic remodeling of MTALTCO1 orthologs

Conservation metrics provide a valuable complement to experimental data, adding due weight to any arguments of hypothesized function (Gruschus et al., 2023). However, ORFs that overlap mitochondrial protein-coding genes pose a direct challenge to standard conservation assessment methods and exhibit unconventional behaviors conservation-wise. MTALTCO1 is found in its full 259 amino acid form only in *Homo sapiens* and *H. heidelbergensis*, with pseudogenization via nonsense mutations observed in closely related Great Apes. This pattern is mirrored by other alternative mitochondrial proteins such as CYTB-187AA and MTALTND4 (Supplementary Material: Supporting Text 3), despite their established causal roles in *Homo sapiens* cellular models. Interestingly, the full-length MTALTCO1 ORF reappears occasionally across vertebrates, reminiscent of the stochastic pseudogenization and reappearance observed for *Humanin*, despite its widely recognized biological importance. In some species, MTALTCO1 orthologs even surpasses MTALTCO1’s length, extending both upstream and downstream, as epitomized by the passerine bird *Sturnus cineraceus*, where it spans roughly 98% of the entire CO1 sequence (Supplementary Material: Supporting Text 3). The mosaic and transient conservation pattern observed in mitochondrial alternative proteins like MTALTCO1 challenges classical models of protein evolution as well as current frameworks in the *de novo* gene birth literature. Although de novo models can accommodate gene fusion and fission events (Andersson et al., 2015), in this context they would require either the persistence of functional domains serving as invariant nucleation points for novel gene constructs, or the emergence of function *ex nihilo*. The former scenario appears incompatible with the rapid and seemingly stochastic remodeling of ORFs observed across primates, while the latter is difficult to reconcile with both the cross-species conservation of molecular features and the lack of positional constraint observed at the population level.

While a fully established conceptual framework linking these noisy conservation signals to biological function is still lacking, there is clear evidence of selection that cautions against excluding conservation from functional hypotheses. By considering evolutionary conservation, our approach supports the idea that MTALTCO1 may play roles in molecular organization, particularly in functions that could tolerate high sequence volatility. This gap nonetheless highlights a critical frontier in the study of mitochondrial alternative proteins, and resorting to ahistorical theories of function to circumvent these challenges might prove epistemically risky (Ardern et al., 2020; Brandon, 2013). Even approaches focused solely on causal roles ultimately rest on implicit historical assumptions about the nature and study of biological function (Garson, 2019), particularly given their foundation in frameworks optimized for canonical proteins, which differ markedly from mtaltORFs.

## CONCLUSION

We provide multiple lines of evidence demonstrating that MTALTCO1 is a protein expressed within mitochondria, incorporating codons that necessitate contextual reassignment, and argue that its biophysical properties and conservation patterns are consistent with scaffolding. This study broadens the conceptual framework for mitochondrial genome expression and advances the understanding of mitochondrial alternative proteins, particularly with respect to their conservation dynamics. Future experimental validation will be essential to establish the biological role of MTALTCO1 and to test the functional hypothesis proposed here. We argue that such efforts should be guided by the specific features of MTALTCO1, its unconventional conservation pattern, physicochemical properties, and interactor profile, to inform the design of appropriately tailored characterization methods.

## CODE AVAILABILITY

The entire code used in generating data for this paper will be made available upon publication in a journal.

