## Supplementary Material for "MTALTCO1: a 259 amino-acid long mtDNA-encoded alternative protein that challenges conventional understandings of mitochondrial genomics" (from another version of this preprint: version 3, 2025-06-25)

#### SUPPORTING TEXT, FIGURES AND TABLES

##### Supporting Text 1: Detailed Material and Methods

###### *Protein extraction and Western Blot*

10-cm petri dishes containing cells were first washed twice with PBS, harvested, and then sonicated at an amplitude of 75% three times for 3 seconds each using a dismembrator sonicator (Fisher) in a homogenization buffer (pH 7.4) containing 10 mM HEPES, 150 mM NaCl and a cocktail of protease inhibitors (benzamidine, PMSF, aprotinin and leupeptin). After sonication, Triton X-100 (10%) was added to achieve a final concentration of 1%, and the samples were incubated for 20 minutes at 4°C with agitation. The samples were then centrifuged at 10,000 rpm for 10 minutes at 4°C, and the supernatant was collected and kept on ice. Total protein concentration was determined using the Bradford assay, and aliquots containing 100 µg of proteins were prepared and stored at -80°C for later use. As mentioned in the main text, various tissue lysates (kidney, heart, testis, brain, liver, skeletal muscle, spleen) were also obtained from Novus Biologicals (NB820-59231; NB820-59217; NB820-59266; NB820-59177; NB820-59232; NB820-59253; NB820-59259).

Samples were suspended in Laemmli Sample Buffer (LSB) 1X containing DTT and ß-mercaptoethanol and heated for 5 minutes at 95°C. They were then loaded onto Tricine-SDS-PAGE 3-part gels (separating, spacer, and compaction gels), ideal for protein separation ranging from 1 to 100 kDa, at an equal protein concentration of 100 µg. The samples were migrated at low voltage (~43 V) overnight at room temperature and subsequently transferred onto a pre-hydrated PVDF membrane with a 0.2 µm pore size for 1 hour at 4°C and 1000 mA in electrophoresis transfer buffer (Towbin buffer). The membrane was then incubated for 1 hour at room temperature in a PBS blocking buffer containing 5% non-fat dry milk and 0.05% Tween-20. Following blocking, the membranes were incubated with primary antibodies for 1.5 hour at room temperature. The primary antibodies used were diluted in PBS with 0.05% Tween-20 as follows: "rabbit anti-MTALCO1" (1:1000) (MediMabs), "mouse anti-ATP5" (1:2000) (Abcam [ab14748]), "mouse anti-CO1" (1:1000) (LifeSpan BioSciences, Inc. [LS-C172110]), "rabbit anti-ND4" (1:1000) (Boster Biological Technology Co., Ltd. [A04180-1]), and "mouse anti-actin" (1:2000) (Abcam [ab1801]). After incubation with primary antibodies, the membranes were washed 3 times for 10 minutes each with TBS-T. They were then incubated with goat anti-rabbit IgG (1:2000) or goat anti-mouse IgG (1:2000) secondary antibodies coupled to horseradish peroxidase (HRP), diluted in PBS-T, for 30 minutes at room temperature. Following incubation, the membranes were washed 3 times for 10 minutes with TBS-T and once for 5 minutes with TBS at room temperature. Equal volumes of chemiluminescent substrate and enhancer solutions West-Pico SuperSignal (Thermo Fisher Scientific). were added to the membranes, and the signals were captured using a FUSION FX chemiluminescence imaging system. To verify the specificity of the anti-MTALTCO1 antibody, the synthetic antigenic peptide was added to the primary antibody solution at a 10X concentration before incubation. This step was performed to competitively chelate every antigenic site of the primary antibody, confirming the attenuation of the specific MTALTCO1 band.

***Immunofluorescence***

Coverslips were incubated in 95% ethanol for 1 hour and then placed in a 24-well PCA detachable microscope slide and covered with a 50µg/mL Poly-D-Lysine (PDL) solution for 2 hours. HeLa cells were cultured on these slides in DMEM supplemented with 10% calf bovine serum, fungizone, penicillin and streptomycin, and maintained at 37˚C with 5% $CO_2$. Each well was washed with PBS and fixed with paraformaldehyde (4%, for 20 minutes), followed by 3 consecutive 5-minutes washes with PBS. To permeabilize the cell membranes, Triton (0.2%) was added for 4 minutes, followed by 3 consecutive 5-minutes washes with PBS. Then, a 30-minutes blocking step at room temperature was performed with PBS supplemented with 1% BSA and 2% NGS. This was followed by a 2-hours incubation with a mixture of 1:50 anti-MTALTCO1 rabbit and 1:250 anti-ATP5 mouse primary antibodies in PBS (1% NGS). After three 5-minutes PBS washes, the cells were incubated with secondary IgG antibodies (respectively Alexa Fluor TM 488 anti-rabbit [1:500] and Alexa Fluor TM 488 anti-mouse [1:500]) in PBS (1% NGS) for 1 hour at room temperature. Finally, the cells were washed three times with PBS for 5 minutes and mounted in DAPI mounting medium (aqueous, fluoroshield) (ab104139), then visualized using a Zeiss-LSM800 confocal microscope.

***Immunoprecipitation and mass spectrometry analysis***

Six 10-cm petri dishes containing HEK-293T cells at a confluence of 80% were washed twice with PBS and harvested by mechanical scraping in a total of 2 mL of lysis buffer containing 10 mM HEPES, 150 mM NaCl, and a cocktail of protease inhibitors (benzamidine, PMSF, aprotinin, and leupeptin). Cell lysis was performed by sonication at an amplitude of 75% three times for 3 seconds each using a dismembrator sonicator. After sonication, Triton X-100 (10%) was added to a final concentration of 1% and samples were incubated for 20 minutes at 4°C with agitation. They were centrifuged at 14 000 rpm for 20 minutes at 4°C and the supernatant was collected and kept on ice.

90 µl of Protein A Mag Sepharose magnetic beads were washed in a wash buffer containing 50mM of tris-buffered saline with 2M urea at a pH of 7.5. To pre-clear the cell supernatant, 2mL of the supernatant collected was added to 20 µl Protein A Mag Sepharose

magnetic beads (Cytiva) and incubated for 1 hour at RT with agitation. 50 uL of previously washed Protein A Mag Sepharose magnetic beads were equilibrated by adding 1000 µl of binding buffer containing 50mM of tris-base and 150mM of NaCl at a pH of 7.5. These were then divided in two different tubes and the supernatant was then removed. 25 µl of rabbit anti-MTALTCO1 antibody (treatment) was added to one of the tubes containing the beads and 25uL of rabbit pre-immune serum (control) was added to the other tube. 500 µl of the lysis buffer was added to avoid the magnetic beads from drying and these were incubated for 20 minutes at RT with agitation.

The supernatants were then removed, and unbound antibodies were removed by adding 500 µl of binding buffer to the samples. The supernatant was removed, and the crosslinking was achieved by first adding 500 µl of 200mM triethanolamine pH 8.9 to the samples. Then, the supernatant was then removed and 500 µl of 200mM triethanolamine with 50mM dimethyl pimelimidate (DMP) pH 8.9 was added to the samples and incubated for 1 hour at RT with agitation. After this incubation, the samples were washed once for 1 minute at RT under agitation with 500 µl of 200mM triethanolamine pH 8.9. The residual crosslinking reagent was blocked by adding 500 µl of 100mM ethanolamine pH 8.9 and incubating the samples for 15 minutes at RT under agitation. The supernatant was then removed, and unbound antibody was removed by incubating the samples for 1 minute at RT under agitation with 500 µl of 0.1M glycine with 2M urea pH 2.9. Finally, the magnetic beads were 1quilibrated by washing them twice for 5 minutes at RT under agitation in 500 µl of binding buffer. The supernatant was removed before 1mL of the pre-cleared cell lysate was added to each sample. The beads and cell lysates were incubated overnight at 4˚C under agitation.

The magnetic beads were washed 6 times with tris-buffered saline with 2M urea and resuspended in 50 µl of 100 mM Tris-HCl pH 8 before being sent to the proteomics core facility of the Institute for Research in Immunology and Cancerology (IRIC, University of Montreal) for MSmass spectrometry analysis. At the IRIC, samples were reconstituted in 50 mM ammonium bicarbonate with 10 mM TCEP [Tris (2-carboxyethyl) phosphine hydrochloride; Thermo Fisher Scientific], and vortexed for 1 h at 37°C. Chloroacetamide (Sigma-Aldrich) was added for alkylation to a final concentration of 55 mM. Samples were vortexed for another hour at 37°C. One microgram of trypsin was added, and digestion was performed for 8 h at 37°C. Samples were dried down and solubilized in 5% ACN-4% formic acid (FA). The samples were loaded on a 1.5ul pre-column (Optimize Technologies, Oregon City, OR). Peptides were separated on a home-made reversed-phase column (150-µm i.d. by 200 mm) with a 56-min gradient from 10 to 30% ACN-0.2% FA and a 600-nl/min flow rate on a Easy nLC-1200 connected to a Exploris 480 (Thermo Fisher Scientific, San Jose, CA). Each full MS spectrum acquired at a resolution of 120,000 was followed by tandem-MS (MS2MS-MS) spectra acquisition on the most abundant multiply charged precursor ions for 3s. MS2Tandem-MS experiments were performed using higher energy collision dissociation (HCD) at a collision energy of 34%. The data were processed using PEAKS X Pro (Bioinformatics Solutions, Waterloo, ON) and a Uniprot database. Mass tolerances on precursor and fragment ions were 10 ppm and 0.01 Da, respectively. Fixed modification was carbamidomethyl (C). Variable selected posttranslational modifications were acetylation (N-ter), oxidation (M), deamidation (NQ), phosphorylation (STY).

The data were visualized with Scaffold 5.0 (protein threshold, 99%, with at least 2 peptides identified and a peptide threshold of 95%). Known contaminants were removed: Keratin, Glutathione S-transferase, Serum Albumin, Trypsin, Tubulin, Decoys and Immunoglobulins. To obtain possible pathways in which MTALTCO1 interacts, possible interactors were identified in both the pulldown and co-immunoprecipitation experiments. To ensure the selection of only the most robust candidate interactors, a stringent filtering pipeline was applied. Proteins were retained if they exhibited a total spectral count of at least three in the treatment condition. Furthermore, the difference in spectral counts between the treatment and control conditions (treatment minus control) had to be equal to or greater than three. Finally, each selected protein was required to have at least two more exclusive peptides identified in the treatment compared to the control. The proteins identified in the co-immunoprecipitation and pull-down experiments were submitted to functional enrichment analysis using the Gene Ontology (GO) and Kyoto Encyclopedia of Genes and Genomes (KEGG) databases.

***Pull down assay and mass spectrometry analysis***

Competent cells (Thermo Scientific BL21(DE3) Competent Cells) were transformed following manufacturer's instructions. Specifically, 1 µl of GST-tagged MTALTCO1 expression plasmid (LifeTein, Somerset, NJ, USA) was added to 50 µl of BL21(D3) competent cells originating from an E. coli B strain (a GST expression plasmid was used as control). Cells were incubated on ice for 30 min, then heat shocked for 30 seconds at 42˚C before returning on ice for 2 min. Then, 250 µl of LB broth growth medium was added to the samples, which were placed in a shaking incubator at 225 rpm for 1 h at 37˚C. Transformed cells were then spread on LB plates containing 25 µl of ampicillin and incubated at 37˚C overnight.

For protein expression, a transformant colony was picked and inoculated in 5 ml of LB medium with 10 µl ampicillin and incubated overnight at 37˚C under continuous agitation, allowing it to grow to saturation. This sample was added to 250 ml of fresh LB medium containing 250 µl of ampicillin and incubated at 37˚C under continuous agitation until an optical density at 600nm of 0.5 was obtained. Then, 250 µl of IPTG was added to the bacterial cultures and incubated at 37 ˚C for 2h. The induced culture was centrifuged at 4200 rpm for 10 min at 4˚C and the pellet was resuspended in 10 ml of PBS 1X with urea 8M. The resuspended pellets were lysed by sonication, using three bursts of 3 seconds at 1-minute intervals at an amplitude of 75%. Triton X100 was added to a final concentration of 1%. The lysed pellets were incubated under agitation at 4 ˚C for 20 minutes before being centrifuged at 4200 rpm for 10mins at 4 ˚C. The supernatant was stored at -80 ˚C until use.

The GST control and GST-MTALTCO1 bacterial lysates prepared as described above were thawed at RT and incubated for 2.5h with 20 µl of a 50% slurry of Pierce glutathione magnetic agarose beads (Thermo Scientific) at RT for pre-clearing. The beads used for the pre-clearing were discarded, and the pre-cleared bacterial supernatant was kept. For every pull-down done,100 µl of glutathione magnetic agarose beads were washed three times with a wash buffer (composed of 10 mM Tris HCl pH 7.5, 150 mM NaCl, 0.5 mM EDTA and 0.4% Triton-X100), before being added to the GST and GST-MTALTCO1 pre-cleared bacterial supernatant andincubated at RT for 3h under constant agitation. The beads were then washed five times with the wash buffer and resuspended in 100 µl of wash buffer. Six 10-cm petri dishes containing HEK-293T cells at a confluence of 80% were washed twice with PBS and harvested by mechanical scraping in a total of 2 mL of lysis buffer containing 10 mM HEPES, 150 mM NaCl, and a cocktail of protease inhibitors (benzamidine, PMSF, aprotinin, and leupeptin). Cell lysis was performed as above for co-immunoprecipitations. 1 mL of the HEK-293T cell lysate was added to 20 µl of beads coupled with GST- MTALTCO1 (treatment) and 1 mL of the HEK-293T cell lysate was added to 3 µl of beads coupled with GST (control). These were incubated overnight at 4˚C under constant agitation. Once the samples were washed 6 times with lysis buffer and resuspended in 50 µl of 100 mM Tris-HCl pH 8, they were sent to the proteomics core facility of the Institute for Research in Immunology and Cancerology (IRIC, University of Montreal) for mass spectrometry analysis. Samples were treated and analyzed as above for immunoprecipitations.

The resulting data were analyzed with Scaffold 5.0, using a 99% protein threshold and 95% peptide threshold. Total spectrum counts and exclusive unique peptide count were determined for each protein detected in both experimental and control samples. Proteins identified in both experiments were filtered and compiled (Supplementary Table below). Filtered proteins were analysed by the software STRING (string-db.org), to determine enriched pathways MTALTCO1 may be involved in (Szklarczyk et al., 2025).

#### Supporting Text 2: Mitochondrial origin of MTALTCO1

##### *Mass-spectrometry peptides, antigenic peptide and NUMTs*

It is presently unclear whether most of mitochondrial-derived peptides (MDPs) are translated in the cytoplasm from exported transcripts using the standard genetic code, or in the mitochondrial matrix using the mitochondrial genetic code, as many could be translated from either code to give identical proteins. The few MDPs identified so far have been assumed to derive from cytoplasmic translation on the basis they contained arginine residues encoded by AGA or AGG codons (usually considered stop codons in the mitochondrial code), or at least neither their mitochondrial translation nor cytoplasmic translation has been rigorously investigated (Gruschus *et al.*, 2023). Notable exceptions include MTALTND4, which was demonstrated to originate from mitochondrial translation (Kienzle *et al.*, 2023), and CYTB187-AA, which is translated in the cytoplasm and subsequently rerouted to mitochondria through a novel targeting signal (Hu *et al.*, 2024).

To establish a mitochondrial origin, one of the first steps involves excluding nuclear pseudogenes as potential confounders in both antigenic peptide recognition and mass spectrometry–derived peptide matches (see NUMTs - Ricchetti *et al.*, 2004). The identification of antigenic peptide sequences and the assessment of peptide-spectrum match specificity rely on background checks against the annotated proteome. Since pseudogene-derived ORFs are typically not included in such annotations but could theoretically generate similar peptides, we systematically evaluated our antigenic peptide and complete peptide matches against both the canonical proteome and pseudogene sequences. The closest sequence similarity among CO1 pseudogenes is observed with MTCO1P12; all other pseudogenes exhibit less than 80% identity at the nucleotide level with the sequence corresponding to MTALTCO1 (for the antigenic peptide sequence), and their putative ORFs are often fragmented (Figure S1A). Our antigenic peptide (PNAPLRLIRPNHSSPTSPISPSPSCWHHYTTN) does not align with any NUMT, as the sequences they hold that match our peptide are surrounded by stop codons and lack cytoplasmic translation initiation sites (Figure S1B). Even under the improbable scenario of translation being initiated from alternative upstream start codons – situated a few codons downstream of the stop codon and upstream of the antigenic peptide – the resulting product would be a truncated protein, far smaller than the ~31 kDa band detected in our western blot analyses. Of our MS-peptides, some had exact sequence matches but were not part of an open reading frame, and others had no sequence match with any pseudogene (Figure S1D).

##### *AGR codons might be reassigned to code for arginine in mitochondrial alternative frames*

MTALTCO1 contains AGR codons (AGA and AGG), which presents a challenge for claims of mitochondrial origin, as these codons are typically interpreted as stop signals in the vertebrate mitochondrial genetic code due to the absence of corresponding tRNAs (Watanabe & Yokobori, 2011). However, in the case of MTALTCO1, these AGR codons do not seem to be bypassed as stop signals but appear to be correctly translated as arginine residues: our findings strongly suggest that AGR codons are being recognized as arginine in the mitochondrial translational context. Further support comes from other putative mitochondrial alternative proteins, such as MTALTND4 and Gau, which also contain AGR codons and are believed to be translated within mitochondria (see source articles for their arguments on the matter: Kienzle *et al.*, 2023 and Faure *et al.*, 2011, respectively). These observations challenge the prevailing assumption that translation in mitochondria invariably terminates at AGR codons, suggesting instead that, under certain conditions, these codons can be reinterpreted as coding for arginine.

It is important to emphasize that we do not suggest AGR codons are generally decoded as arginine in canonical reading frames. On the contrary, their strict avoidance in reference-frames suggests strong selective pressure against their usage, which would be difficult to rationalize if AGR codons were routinely decoded as arginine.

##### *Putative mechanisms of an overlapping genetic code*

The mechanism by which AGA and AGG codons are interpreted as arginine in alternative mitochondrial reading frames remains unresolved. Several hypotheses may account for this phenomenon. One possibility involves the import of cytosolic tRNAs capable of recognizing AGR codons. Cytosolic tRNAs charged with arginine bear AGR anticodons, and mitochondria are known to import cytosolic tRNAs (Rubio *et al*., 2008). If these tRNAs are indeed present and functional in the mitochondrial matrix, they could enable translation of AGR codons as arginine under certain circumstances, thereby supporting the expression of overlapping genes that would otherwise be interrupted by premature stop signals.

Another hypothesis involves mitochondrial suppressor tRNAs, particularly antisense antitermination tRNAs, as proposed by Seligmann (2011, 2012). These tRNAs possess anticodons complementary to stop codons, potentially enabling stop codon reassignment if aminoacylated with arginine residues. Such a mechanism may facilitate "overprinting", i.e. the expression of additional proteins from overlapping reading frames otherwise constrained by stop codons. Supporting this model, co-evolutionary analyses have revealed a positive correlation between both AGR codon usage and overlapping genes in frame +3 (the frame of MTALTCO1) with the presence of corresponding antisense antitermination tRNAs across 57 primate mitochondrial genomes (Seligmann, 2011). Preliminary evidence suggests that *Homo Sapiens* may lack antitermination tRNAs matching AGR codons and rather rely on the import of tRNAs (Seligmann, 2011). However, these conclusions are based on early secondary structure prediction tools (Lowe & Eddy, 1997), and more recent studies have since identified novel cloverleaf tRNA structures (Mukai *et al.,* 2017), warranting further investigation.

While these two scenarios could account for the translation of mitochondrial genes containing AGR codons, they do not explain the strict avoidance of these codons in reference genes, unless such mechanisms are tightly regulated and restricted to certain conditions. This would imply that the expression of MTALTCO1 and other putative mitochondrial ORFs containing AGR codons depends on context-specific factors, such as the regulated import of tRNAs or the conditional transcription of antisense tRNAs. As pointed out by Seligmann (2012), these regulatory determinants may define the circumstances under which alternative mitochondrial reading frames can be expressed.

A third, more speculative possibility involves ribosomal-level mechanisms, such as modified ribosomes or altered decoding dynamics (e.g., ribosome stalling, release and rescue), which could shift or relax codon recognition specificity. AGA and AGG are not conventional stop codons, and proper termination at AGR sites in canonical reading frames requires the participation of specific release factors (Kruger *et al*., 2023). These processes may operate differently in alternative reading frames, potentially allowing context-dependent misrecognition of AGR stop codons. While such mechanisms might account for translational readthrough events, they do not, by themselves, constitute true codon reassignment. However, in conjunction with other processes, such as those listed above, they could contribute to the translation of AGR-containing sequences in alternative frames like that of MTALTCO1.

Any plausible model of AGR codon reassignment in mitochondria must reconcile two key observations: the apparent ability of alternative reading frames to decode AGR as arginine without readthroughs, and the persistent avoidance of AGR codons in canonical reference frames, where they function as efficient termination signals. While the decoding of AGR codons as arginine in alternative frames is experimentally supported by a suite of evidence, the underlying mechanism remains speculative.

##### *MDPs challenge conventional models of mitochondrial expression*

We have robustly established the existence of MTALTCO1 through multiple independent lines of evidence, supporting the notion that alternative proteins may exhibit unique expression dynamics. Similarly, other MDPs challenge conventional understandings of mitochondrial expression dynamics, albeit in different ways. For example, the SHMOOSE ORF (Miller *et al*., 2023) overlaps tRNA Serine and ND5 genes, suggesting for the first time that either translation might occur prior to the excision of tRNAs from the full-length mRNA transcript, an early step in the maturation of mitochondrial mRNAs, or that variable cleavage of the full length transcript exports mRNAs comprising of coding and non-coding regions (Rackham & Filipovska, 2022). CYTB-187AA represents a novel case of mitochondrial mRNA export followed by cytoplasmic translation and subsequent mitochondrial re-entry via a novel mitochondrial targeting sequence (Hu *et al*., 2024). Recent mitochondrial ribosome profiling studies have also suggested that non-canonical start codons might initiate translation in alternative reading frames (Wakigawa *et al*., 2023).

Since even recent models of mitochondrial expression largely overlook mtaltORFs, they cannot be expected to fully capture their possibly unique expression dynamics. As such, investigating the mitogenome for altORFs may benefit from approaches that remain open to outcomes beyond established models.

##### *References*

Gruschus, J. M., Morris, D. L., & Tjandra, N. (2023). Evidence of natural selection in the mitochondrial-derived peptides humanin and SHLP6. *Scientific Reports*, *13*(1), 14110.

Kienzle, L., Bettinazzi, S., Choquette, T., Brunet, M., Khorami, H. H., Jacques, J. F., ... & Breton, S. (2023). A small protein coded within the mitochondrial canonical gene nd4 regulates mitochondrial bioenergetics. *Bmc Biology*, *21*(1), 111.

Hu, Z., Yang, L., Zhang, M., Tang, H., Huang, Y., Su, Y., ... & Liu, X. (2024). A novel protein CYTB-187AA encoded by the mitochondrial gene CYTB modulates mammalian early development. *Cell Metabolism*.

Ricchetti, M., Tekaia, F., & Dujon, B. (2004). Continued colonization of the human genome by mitochondrial DNA. *PLoS biology*, *2*(9), e273.

Watanabe, K., & Yokobori, S. I. (2011). tRNA modification and genetic code variations in animal mitochondria. *Journal of nucleic acids*, *2011*(1), 623095.

Faure, E., Delaye, L., Tribolo, S., Levasseur, A., Seligmann, H., & Barthélémy, R. M. (2011). Probable presence of an ubiquitous cryptic mitochondrial gene on the antisense strand of the cytochrome oxidase I gene. *Biology direct*, *6*, 1-22.

Rubio, M. A. T., Rinehart, J. J., Krett, B., Duvezin-Caubet, S., Reichert, A. S., Söll, D., & Alfonzo, J. D. (2008). Mammalian mitochondria have the innate ability to import tRNAs by a mechanism distinct from protein import. *Proceedings of the National Academy of Sciences*, *105*(27), 9186-9191.

Seligmann, H. (2011). Two genetic codes, one genome: frameshifted primate mitochondrial genes code for additional proteins in presence of antisense antitermination tRNAs. *Biosystems*, *105*(3), 271-285.

Seligmann, H. (2012). An overlapping genetic code for frameshifted overlapping genes in Drosophila mitochondria: antisense antitermination tRNAs UAR insert serine. *Journal of Theoretical Biology*, *298*, 51-76.

Lowe, T. M., & Eddy, S. R. (1997). tRNAscan-SE: a program for improved detection of transfer RNA genes in genomic sequence. *Nucleic acids research*, *25*(5), 955-964.

Mukai, T., Vargas-Rodriguez, O., Englert, M., Tripp, H. J., Ivanova, N. N., Rubin, E. M., ... & Söll, D. (2017). Transfer RNAs with novel cloverleaf structures. *Nucleic Acids Research*, *45*(5), 2776-2785.

Krüger, A., Remes, C., Shiriaev, D. I., Liu, Y., Spåhr, H., Wibom, R., ... & Rorbach, J. (2023). Human mitochondria require mtRF1 for translation termination at non-canonical stop codons. *Nature communications*, *14*(1), 30.

Miller, B., Kim, S. J., Mehta, H. H., Cao, K., Kumagai, H., Thumaty, N., ... & Alzheimer's Disease Neuroimaging Initiative*. (2023). Mitochondrial DNA variation in Alzheimer's disease reveals a unique microprotein called SHMOOSE. *Molecular psychiatry*, *28*(4), 1813-1826.

Rackham, O., & Filipovska, A. (2022). Organization and expression of the mammalian mitochondrial genome. *Nature Reviews Genetics*, *23*(10), 606-623.

Wakigawa, T., Mito, M., Yamashiro, H., Tomuro, K., Tani, H., Tomizawa, K., ... & Iwasaki, S. (2023). The complexity and dynamics of in organello translation assessed by high-resolution mitochondrial ribosome profiling. *BioRxiv*, 2023-07.

#### Supporting Text 3: Conservation of MTALTCO1

As mentioned in the main text, the conservation of MDPs Overlapping Protein-Coding Genes (MOPCG) has not been investigated beyond pseudogenization assays. Consequently, there are no established benchmarks linking conservation patterns to function in MOPCGs. As the nature, scale, and intensity of selection acting on these genes remain largely unknown, we aimed to explore a wide range of conservation signals. This approach was intended to minimize *a priori* bias regarding which features are most informative in MOPCGs.

##### *Frequent fusion and fission events of MTALTCO1's open reading frame*

MTALTCO1's ORF is pseudogenized in several closely related primates, initially leading us to suspect that it might be a lineage-specific *de novo* gene with limited evolutionary conservation. However, further analysis of the fusion and fission dynamics of the MTALTCO1 ORF across vertebrates revealed a surprising pattern: the ORF not only re-emerges in multiple lineages but, in some cases, expands beyond its original boundaries. Notably, in the passerine bird *Sturnus cineraceus*, the ORF extends across nearly the entire CO1 gene (Figure S2).

To evaluate whether such volatility is unique to MTALTCO1, we examined the persistence of CYTB187-AA, MTALTND4 and MTALTCO1 orthologs in Great Apes. We found that these sequences, too, undergo dynamic remodeling even among closely related species (see Figure S3). Notably, the CYTB187-AA ORF contains a mitochondrial targeting sequence spanning amino acids 1–37, a feature thought to underlie the protein's causal role. This segment is separated from the bulk of CYTB187-AA's ORF in all other Great Apes, and is not translated in *Pongo pygmaeus* and other primates. While theoretical models of *de novo* gene birth can accommodate ORF fusion and fission events, the highly stochastic and rapid remodeling observed here, especially for MTALTCO1, is difficult to reconcile with existing frameworks. Complicating interpretation further, vertebrate mitochondrial genomes have traditionally been considered largely evolutionarily static (Shtolz & Mishmar, 2023) and to date, no models exist to describe mitochondrial *de novo* gene origination, at least in vertebrates. Taken together, these observations led us to reconsider our initial assumption and to explore the conservation of MTALTCO1, despite its apparent volatility.

***Challenges in studying the conservation of overlapping mtaltORFs***

Evidence of selection, whether at the nucleotide level through base-pair conservation or at the amino acid level via purifying or positive selection, is used to probe non-coding regions for unannotated ORFs or to argue for their role in normal cell function (Gruschus *et al*., 2023). These approaches must be carefully adapted when applied to overlapping ORFs, as selection signals from one reading frame can confound interpretation in the other. For instance, a standard method such as PhyloP, can be adapted to detect evolutionary signatures, as was the case for the nuclear alternative protein alt-FUS, whose coding sequence is entirely embedded within the canonical FUS gene (Brunet *et al*., 2021).

Rather than interpreting high conservation as evidence of selection, researchers observed that the region of FUS overlapping with alt-FUS was less conserved than adjacent regions, suggesting evolutionary compromise due to dual-coding constraints. Additional approaches, such as assessing synonymous codon bias, can also be extended to overlapping ORFs by comparing all combinations of synonymous and nonsynonymous substitutions across frame pairings (OLGenie: Nelson *et al*., 2020).

These methods require careful contextualization of the sequences being analyzed, as their interpretability depends heavily on the relative, and often *a priori* unknown, functional importance of the overlapping ORFs. Evolutionary signals detected in overlapping regions are meaningful only when both ORFs exert selective pressures of comparable magnitude. For instance, in the case of alt-FUS, signs of evolutionary compromise could emerge only because both alt-FUS and FUS are likely to contribute similarly in importance to cellular function. If, by contrast, one ORF is overwhelmingly more functionally critical than the other, natural selection will predominantly preserve the more important reading frame, potentially masking or nullifying some evolutionary signatures of the less important one. This is the case in mtDNA, which undoubtedly prioritizes the conservation of reference proteins, and especially so in the context of CO1 which is one of the most conserved mitochondrial genes. In our context, where neighbour divergence in MTALTCO1 is roughly 15 times that of CO1, these approaches offer limited insight.

First, our assessment of nucleotide conservation using FFT-smoothed PhyloP scores sourced from UCSC's 30 primates showed no evidence of dual-coding compromise (Figure S4). We then considered looking at overlapping synonymous bias using the OLGenie framework. However, the extreme asymmetry in amino acid conservation between CO1 and MTALTCO1 in primates likewise limits the informativeness of this approach. In the OLGenie framework, $d_{NN}/d_{NS}$ or $d_{SN}/d_{SS}$ can be used to assess synonymous bias in overlapping frames, and either can be used on a case by case basis, depending on the number of mutations in either gene. Here, the first letter represents the mutation effect (N = nonsynonymous, S = synonymous) in the reference gene, and the second letter represents the effect in the overlapping gene (NN, nonsynonymous/nonsynonymous; SN, synonymous/nonsynonymous; NS, nonsynonymous/synonymous; SS, synonymous/synonymous - refer the OLGenie article for more information). In the case of MTALTCO1, both metrics hold as numerator and/or as denominator pairs of mutations that happen either very frequently, or very little across primates (NN, very rare; SN, very frequent; NS, very rare; SS, very rare). As noted in the OLGenie article, pairs that happen very rarely make for poor estimators of synonymous bias, and we therefore decided to abandon this analysis.

Instead, we hypothesized that evidence for positional conservation might be more readily detectable through position-specific conservation, particularly at codon wobble positions, which could reveal selective pressure on the overlapping reading frame. To test this, we decomposed PhyloP conservation scores by codon position (first, second, and third nucleotide) within the CO1 frame, as done previously by Samandi *et al.*, (2017). The latter showed that nuclear altORFs overlapping annotated protein coding sequences showcase extreme wobble nucleotide phyloP values, being both more variable and more conserved than expected when compared to non-overlapping regions. MOPCGs remain a newly recognized class of proteins, and robust negative controls (i.e., regions known to lack MOPCGs) are currently unavailable. Given the compactness of the mitochondrial genome and the evolutionary incentives for overprinting, the full extent of the mitochondrial alternative proteome remains unknown but probably underestimated. As a result, high wobble nucleotide variability in putative MOPCG regions lacks a meaningful baseline for interpretation. Therefore, in this context, only unusually high conservation at wobble positions, comparable to that seen at first and second codon positions, can be considered informative.

As expected, the first and second codon positions –typically subject to strong amino acid-level constraint– showed the highest conservation (Figure S5). Third (wobble) positions, which are generally more variable, exhibited heterogeneous conservation patterns. Notably, a subset of wobble positions showed elevated conservation levels, comparable to first and second positions, suggesting possible site-specific constraints consistent with dual coding. Smoothing of the PhyloP signals (window size = 25) revealed no obvious regions of high constraint; instead, conserved residues were dispersed throughout the MTALTCO1 reading frame. A sliding-window graphical interface mapping per-nucleotide PhyloP scores onto the MTALTCO1 amino acid sequence is available on our GitHub repository in the conservation.ipynb file, to be published upon publication. Whether these conserved wobble positions reflect functional constraints, RNA structural features, or other regulatory signals relevant to MTALTCO1 remains to be determined and warrants further investigation.

***Moving beyond positional conservation, are molecular features conserved?***

Although there is some evidence of positional conservation for a few residues, the overwhelming majority of residues are quite poorly conserved, as also suggested by elevated primate-level Hamming distances (Figure 3, main text). Various disorder and structure prediction tools suggest MTALTCO1 might be an intrinsically disordered protein (IDP), and we sought to monitor whether molecular features might show evidence of conservation, as IDPs are known to retain molecular features and not positional information (Zarin *et al.*, 2019). Molecular features important to IDPs, as identified by Zarin *et al*., (2019) were retained if they could produce a single-value output per sequence.

The strong asymmetry that favors positional conservation of the CO1 gene reinforces the rationale behind leveraging degeneracy to monitor the unlikeliness of proteins in alternative frames. This approach has been used in viruses to compare lengths of actual ORFs to those of simulated, degenerate ORFs (Schlub *et al*., 2018) and was also used in the Gau paper to show that actual sequences are more self-similar than generated ones (Faure *et al*., 2011). The argument behind this approach is that any deviation from random expectation is a telltale sign of selection. So instead of monitoring the conservation of molecular features throughout primates, as this conservation might only be the outcome of constraints imposed by the CO1 gene, we monitored deviation from random expectation (of degenerate sequences) for these molecular features throughout primates (see results in main text).

In our implementation, MTALTCO1 variants from 27 primate species (full list below) were projected into a KDE landscape generated from 600 degenerate sequences in the +3 reading frame. The goal was to generate a distribution of plausible off-frame ORFs under neutrality, against which real sequences could be ranked. Our approach has some limitations, the first of these being that we obtain a rank, not a conventional p-value. While the interpretation of these is somewhat similar, they do not rely on the same formalism, and to make this distinction clear we ranked sequences from 0 being the most KDE-dense sequence to 1 being the most unlikely sequence (least KDE-dense), thereby avoiding confusion with p-values, wherein low value suggest departures from a given distribution.

We made the choice to rank real MTALTCO1 sequences in the KDE space of simulated degenerate sequences mainly because this is a non-parametric approach that does not assume any underlying distribution to derive p-values. The rationale for a non-parametric approach, that is to avoid assumptions about the underlying density distribution, particularly given the increased granularity and radial asymmetry away from the distribution's center, might seem overly zealous at first glance given the rough radial asymmetry. We trace this decision to our exploratory findings, where departures from neutral expectation were studied to compare MTALTCO1 with every other putative ORF in the +3 reading frame for other mitochondrial protein coding genes (Figure S6). This KDE distribution was highly radially asymmetrical, and p-value formalisms (variably based on some form or combination of Gaussian distributions) were likely to assume some misguided form of distribution. Furthermore, combinatorial constraints meant that some features were quantized, and not on continuous gradients, as can be clearly seen for the length of generated ORFs in figure 3A of the main text. Such 'islands' of probabilities would therefore be strongly inaccurate in their p-value estimates, were we to smooth them in some assumed distribution. In all exploratory and presented results, we sought to retain this formalism to allow for direct comparisons.

Therefore, we considered that generating many degenerate sequences to obtain a precise KDE distribution would be a prudent way forward, yet this approach entails a trade-off. Generating more DSS sequences improves KDE resolution but introduces difficulty for PCA to orient meaningful principal components if the background noise overwhelms real sequence variance. Conversely, too few DSS sequences makes the KDE less stable, increasing noise in the rank estimation. We optimized this balance empirically by examining whether principal components still captured independent features such as amino acid usage. One-to-one comparisons of amino acid usage (example in Figure S7) show that deviations in amino acid usage among real sequences are reliably captured by PCA (Figure 3, main text), provided that the number of simulated sequences is not excessively large. To balance KDE resolution with PCA interpretability, we selected a DSS-to-real sequence ratio that maximized KDE estimation while still allowing PCA to reflect meaningful biological variation.

Full 27 primate list (sequence corresponding in indices to MTALTCO1):

0: NC_012920.1 Homo sapiens mitochondrion
1: NC_001643.1 Pan troglodytes mitochondrion
2: NC_001646.1 Pongo pygmaeus mitochondrion
3: NC_002082.1 Hylobates lar mitochondrion
4: NC_023971.1 Trachypithecus cristatus mitochondrion
5: NC_006901.1 Colobus guereza mitochondrion
6: NC_023961.1 Cercopithecus mitis mitochondrion
7: NC_021947.1 Erythrocebus patas mitochondrion
8: NC_007009.1 Chlorocebus aethiops mitochondrion
9: NC_021956.1 Mandrillus sphinx mitochondrion
10: NC_012670.1 Macaca fascicularis mitochondrion
11: NC_019802.1 Theropithecus gelada mitochondrion
12: NC_001992.1 Papio hamadryas mitochondrion
13: NC_050682.1 Callithrix aurita mitochondrion
14: NC_021960.1 Saguinus oedipus mitochondrion
15: NC_064162.1 Cacajao calvus ucayalii isolate FES101 voucher IDSM03679 mitochondrion
16: NC_024630.1 Callicebus lugens mitochondrion
17: NC_018116.1 Aotus nancymaae mitochondrion
18: NC_021951.1 Lagothrix lagotricha mitochondrion
19: NC_012775.1 Saimiri sciureus mitochondrion
20: NC_027825.1 Alouatta seniculus mitochondrion
21: NC_002763.1 Cebus albifrons mitochondrion
22: NC_012762.1 Otolemur crassicaudatus mitochondrion
23: NC_010299.1 Daubentonia madagascariensis mitochondrion
24: NC_010300.1 Eulemur mongoz mitochondrion
25: NC_028210.1 Propithecus verreauxi mitochondrion
26: NC_002765.1 Nycticebus coucang mitochondrion

#### ***References***

Shtolz, N., & Mishmar, D. (2023). The metazoan landscape of mitochondrial DNA gene order and content is shaped by selection and affects mitochondrial transcription. *Communications Biology*, *6*(1), 93.

Gruschus, J. M., Morris, D. L., & Tjandra, N. (2023). Evidence of natural selection in the mitochondrial-derived peptides humanin and SHLP6. *Scientific Reports*, *13*(1), 14110.

Brunet, M. A., Jacques, J. F., Nassari, S., Tyzack, G. E., McGoldrick, P., Zinman, L., ... & Roucou, X. (2021). The FUS gene is dual‐coding with both proteins contributing to FUS‐mediated toxicity. *EMBO reports*, *22*(1), e50640.

Nelson, C. W., Ardern, Z., & Wei, X. (2020). OLGenie: estimating natural selection to predict functional overlapping genes. *Molecular Biology and Evolution*, *37*(8), 2440-2449.

Samandi, S., Roy, A. V., Delcourt, V., Lucier, J. F., Gagnon, J., Beaudoin, M. C., ... & Roucou, X. (2017). Deep transcriptome annotation enables the discovery and functional characterization of cryptic small proteins. *Elife*, *6*, e27860.

Zarin, T., Strome, B., Nguyen Ba, A. N., Alberti, S., Forman-Kay, J. D., & Moses, A. M. (2019). Proteome-wide signatures of function in highly diverged intrinsically disordered regions. *Elife*, *8*, e46883.

Schlub, T. E., Buchmann, J. P., & Holmes, E. C. (2018). A simple method to detect candidate overlapping genes in viruses using single genome sequences. *Molecular Biology and Evolution*, *35*(10), 2572-2581.

Faure, E., Delaye, L., Tribolo, S., Levasseur, A., Seligmann, H., & Barthélémy, R. M. (2011). Probable presence of an ubiquitous cryptic mitochondrial gene on the antisense strand of the cytochrome oxidase I gene. *Biology direct*, *6*, 1-22.

#### SUPPORTING FIGURES

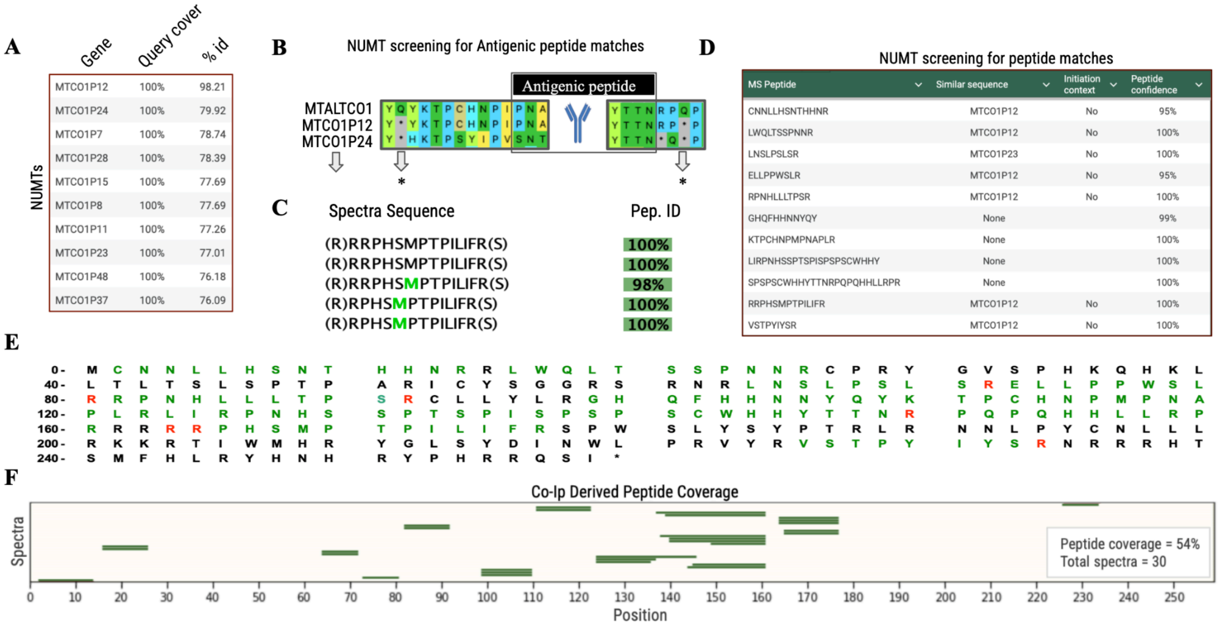

**Figure S1. Global portrait of MTALTCO1: origin, localization, structure and disorder predictions.** (A-B) Pseudogene hits for our antigenic peptide sequence (query), %id is at the nucleotide level. The two best hits are shown in B, with other pseudogenes being increasingly unlikely confounding factors. (C) Spectra with both ATA encoded M and AGR encoded R residues, with corresponding peptide identification probability. (D) Our MS-peptides were searched against translated pseudogenes, with 'similar sequence' being exact sequence matches for these peptides. Initiation context denotes the presence of an upstream ATG encoded Methionine. (A) The amino acid sequence coverage for MTALTCO1 is 54%, for 30 total spectra, 25 exclusive unique spectra and 20 exclusive unique peptides. The amino acids in green and red are present in at least one spectra, with the latter being arginine residues coded by AGA/AGG. (F) Spectral coverage (every green bar represents a spectrum).

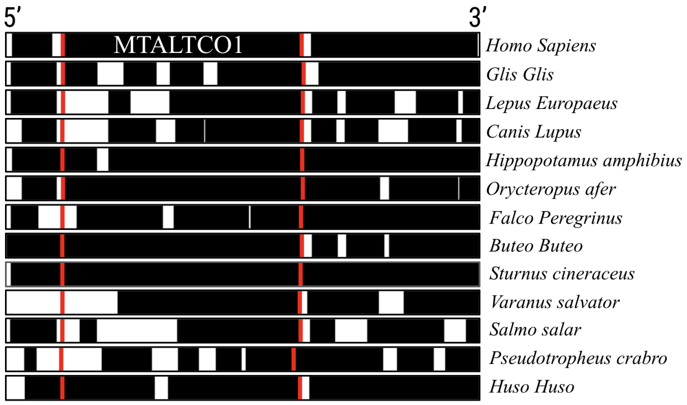

**Figure S2. Dynamic remodelling of MTALTCO1 in a few vertebrates.**
The horizontal bars represent the entire coding sequence of CO1, spanning from nucleotide 0 to 1541 in humans, from left (5') to right (3'). White sections of the bars indicate regions not translated in the +3 reading frame for lack of a start codon, while black sections represent regions between a start and stop codon in the +3 reading frame. MTALTCO1's start and stop codons are marked in red to show the corresponding regions in other vertebrates.

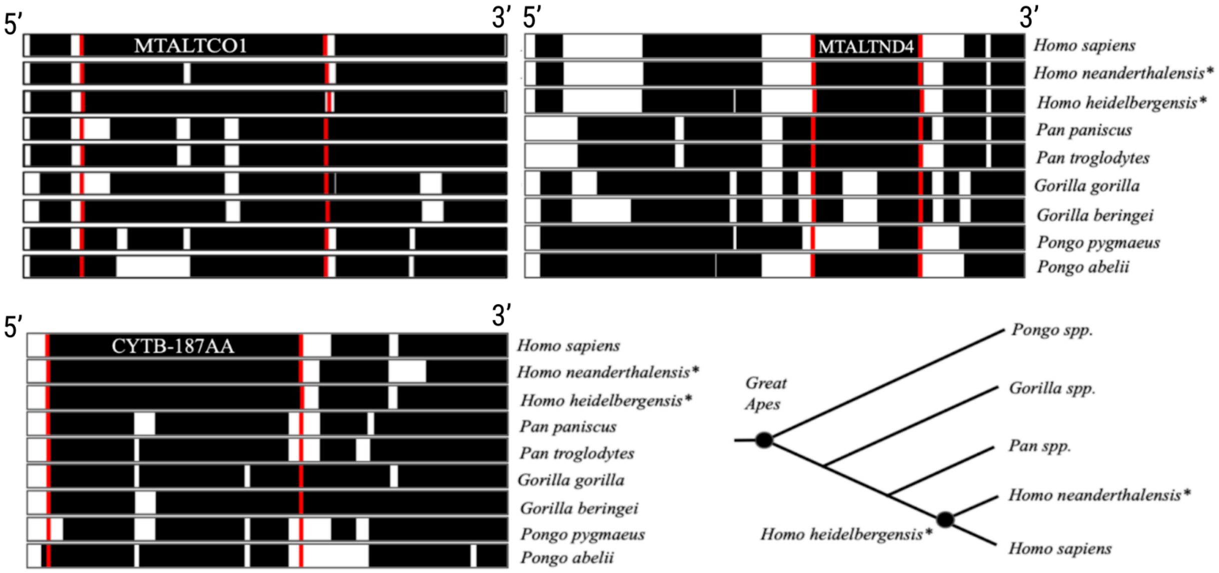

**Figure S3. Dynamic remodelling of mitochondrial alternative ORFs in Great Apes**
Horizontal bars represent the same thing as in figure S2. MTALTCO1's start and stop codons are marked in red to show the corresponding regions in other hominids. Species are arranged top to bottom in an approximately proximal-to-distal order, starting with humans, with the actual phylogeny displayed on the right. Extinct species are marked with an asterisk.

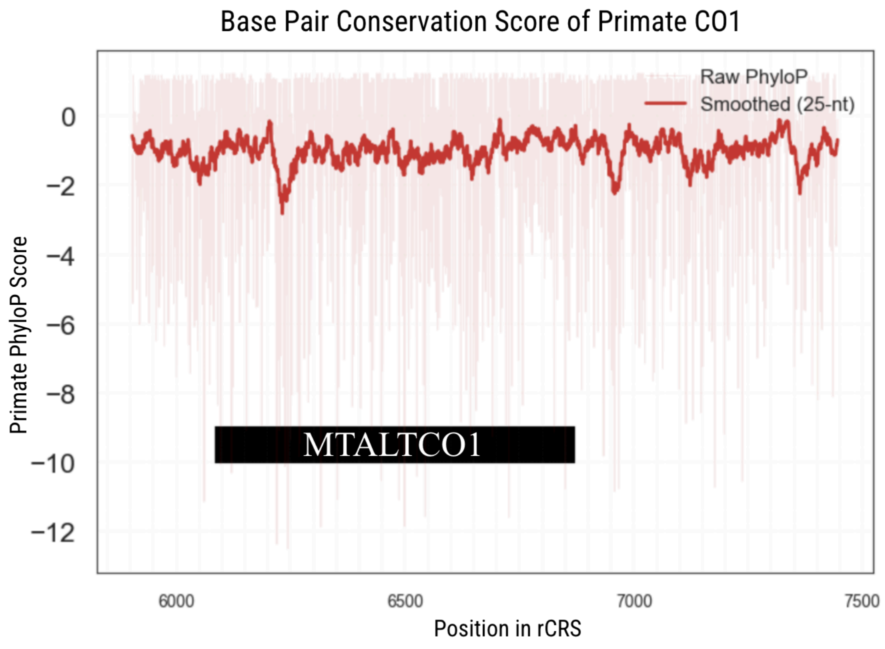

#### Figure S4. Base pair conservation scores for primate CO1

Phylo-P scores of CO1 in 30 primates smoothed over a 25-nucleotide window. Position is with respect to the mitochondrial revised Cambridge Reference Sequence (rCRS).

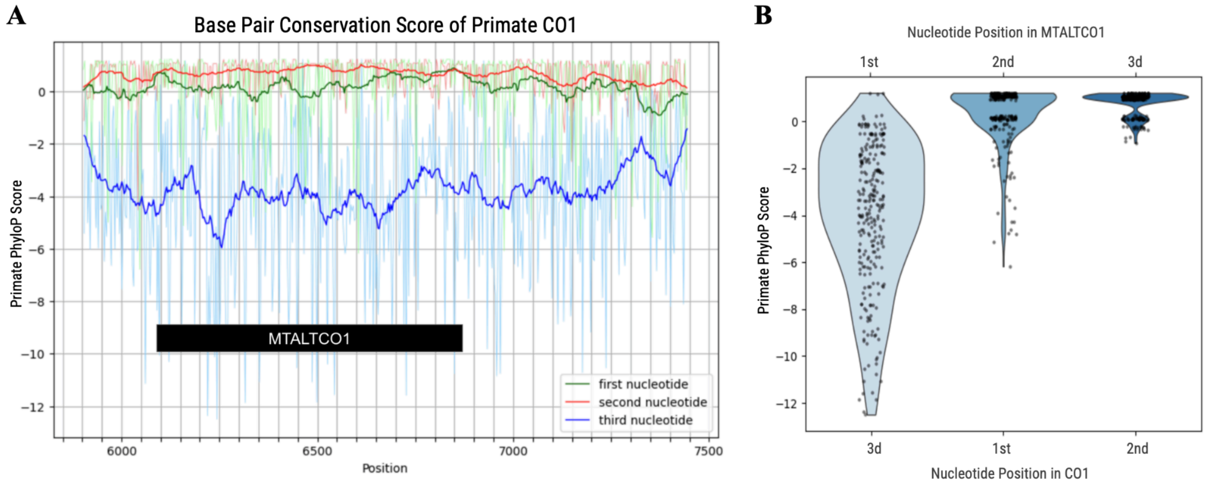

#### Figure S5. Decomposed PhyloP scores for primate CO1 and MTALTCO1

(A) Decomposed Phylo-P scores of CO1 in 30 primates smoothed with over a 25-nucleotide window, with the MTALTCO1 ORF positioned in black. (B) Decomposed PhyloP scores, ordered according to the RF of either CO1 or MTALTCO1.

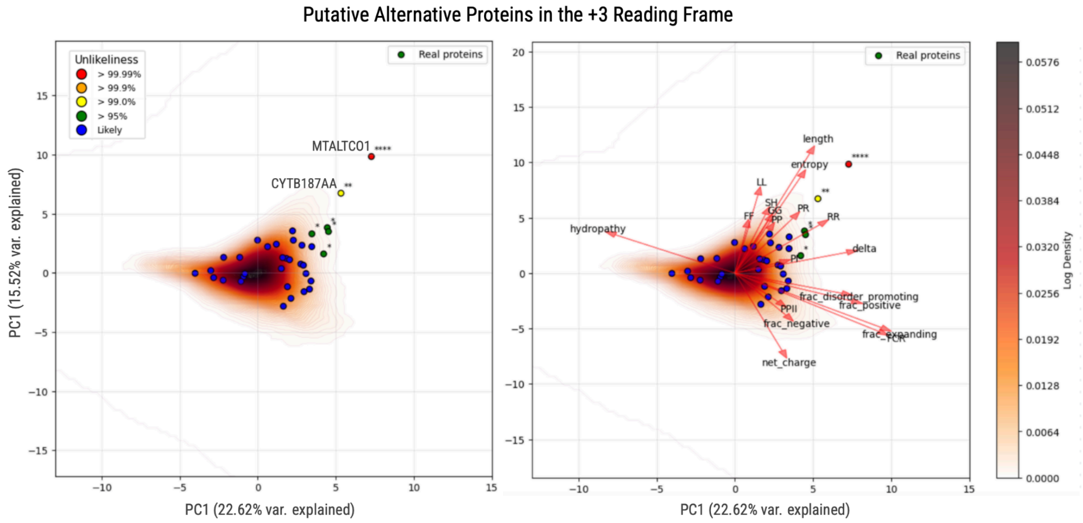

### Figure S6. Exploratory findings with radial asymmetry

Distribution of amino acid usage and molecular features in reduced feature space for generated (topographic contour lines - color-coded by the Log Density) and real protein sequences (scatter - ranked and color-coded by their density percentile) in the +3 reading frame overlapping every mitochondrial protein coding gene. Outliers with experimental confirmation are labelled.

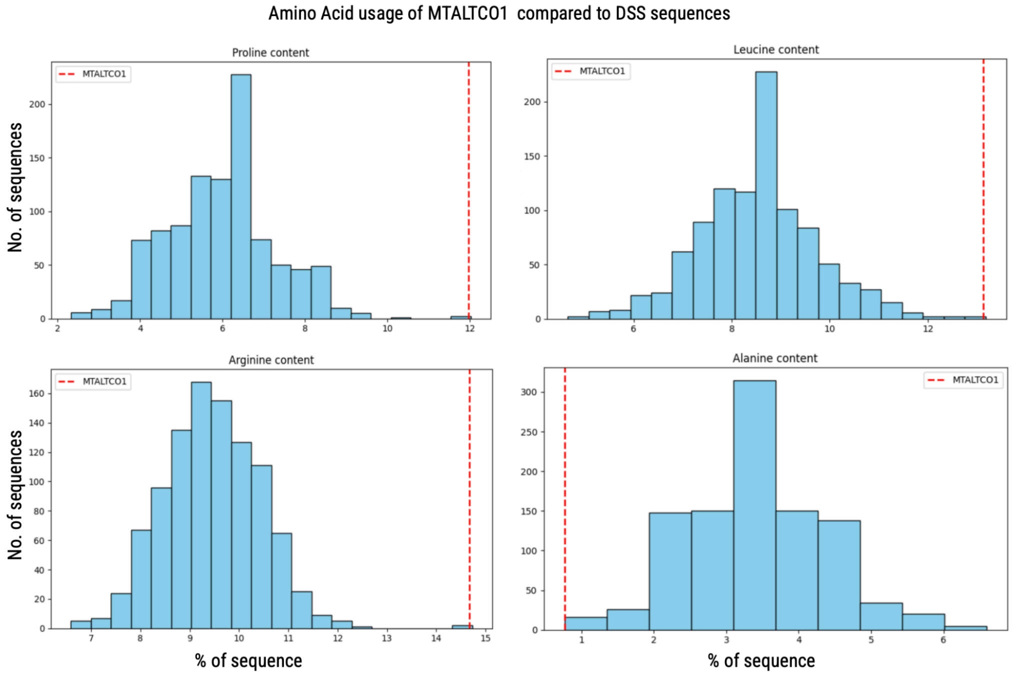

### Figure S7. Amino acid usage of generated sequences compared with MTALTCO1

Four metrics of MTALTCO1 are shown compared to those of proteins translated from the +3 reading frames of generated nucleotide sequences that preserve CO1's sequence. The histogram bars represent the values from generated sequences, while the observed value for MTALTCO1 is marked in red.

#### SUPPORTING TABLES

**Description**

The following three tables present proteins retained as candidate interactors of MTALTCO1 in experiments performed on HEK293T cells. Table 1 includes proteins detected in both the co-immunoprecipitation and GST-pulldown assays, while Tables 2 and 3 list proteins uniquely identified in the co-immunoprecipitation and GST-pulldown experiments, respectively.

To be retained as a potential interactor, a protein had to meet all of the following criteria in the treatment condition (TRMT):

1. A minimum total spectral count of 3;
2. A difference of at least three spectral counts between treatment and control conditions (TRMT – CTRL ≥ 3);
3. At least two additional exclusive unique peptides compared to the control.

In total, 10 proteins were identified in both experimental setups, 60 proteins were exclusive to co-immunoprecipitation, and 227 were exclusive to GST-pulldown.

**Table List**

Table S1. Proteins identified as potential MTALTCO1 interactors in both co-immunoprecipitation and GST-pulldown experiments

Table S2. Proteins identified exclusively in the co-immunoprecipitation experiment

Table S3. Proteins identified exclusively in the GST-pulldown experiment

##### Table S1. Proteins identified as potential MTALTCO1 interactors in both co-immunoprecipitation and GST-pulldown experiments.

| | | | Total Spectrum Count Co-IP (HEK293T) | | | Exclusive Unique Peptide Count Co-IP (HEK293T) | | | Total Spectrum Count GST-pulldown (HEK293T) | | |
|---|---|---|---|---|---|---|---|---|---|---|---|
| Identified Proteins | Accession Number | Molecular Weight | CTRL | TRMT | TRMT-CTRL | CTRL | TRMT | TRMT-CTRL | CTRL | TRMT | TRMT-CTRL |
| Poly [ADP-ribose] polymerase 1 OS=Homo sapiens OX=9606 GN=PARP1 PE=1 SV=4 | P09874 | 113 kDa | 1 | 7 | 6 | 1 | 6 | 5 | 18 | 158 | 140 |
| Desmoplakin OS=Homo sapiens OX=9606 GN=DSP PE=1 SV=3 | P15924 | 332 kDa | 17 | 38 | 21 | 17 | 37 | 20 | 22 | 41 | 19 |
| Junction plakoglobin OS=Homo sapiens OX=9606 GN=JUP PE=1 SV=3 | P14923 | 82 kDa | 5 | 16 | 11 | 5 | 14 | 9 | 11 | 18 | 7 |

| | | | | | | | | | | |
|---|---|---|---|---|---|---|---|---|---|---|
| Filaggrin OS=Homo sapiens OX=9606 GN=FLG PE=1 SV=3 | P20930 | 435 kDa | 10 | 15 | 5 | 10 | 12 | 2 | 1 | 14 | 13 |
| Suprabasin OS=Homo sapiens OX=9606 GN=SBSN PE=1 SV=2 | Q6UWP8 | 61 kDa | 9 | 12 | 3 | 7 | 9 | 2 | 2 | 10 | 8 |
| 60S ribosomal protein L15 OS=Homo sapiens OX=9606 GN=RPL15 PE=1 SV=2 | P61313 | 24 kDa | 2 | 6 | 4 | 2 | 4 | 2 | 5 | 10 | 5 |
| Lysozyme C OS=Homo sapiens OX=9606 GN=LYZ PE=1 SV=1 | P61626 | 17 kDa | 5 | 11 | 6 | 5 | 10 | 5 | 0 | 4 | 4 |
| RuvB-like 2 OS=Homo sapiens OX=9606 GN=RUVBL2 PE=1 SV=3 | Q9Y230 | 51 kDa | 2 | 6 | 4 | 2 | 6 | 4 | 2 | 9 | 7 |
| Retinitis pigmentosa 9 protein OS=Homo sapiens OX=9606 GN=RP9 PE=1 SV=2 | Q8TA86 | 26 kDa | 6 | 10 | 4 | 6 | 8 | 2 | 0 | 3 | 3 |
| 60S ribosomal protein L32 OS=Homo sapiens OX=9606 GN=RPL32 PE=1 SV=2 | P62910 | 16 kDa | 2 | 5 | 3 | 2 | 4 | 2 | 2 | 5 | 3 |

**Table S2. Proteins identified exclusively in the co-immunoprecipitation experiment.**

| | | | Total Spectrum Count Co-IP (HEK293T) | | | Exclusi |
|---|---|---|---|---|---|---|
| Identified Proteins | Accession Number | Molecular Weight | CTRL | TRMT | TRMT-CTRL | CTRL |
| Thyroid receptor-interacting protein 11 OS=Homo sapiens OX=9606 GN=TRIP11 PE=1 SV=3 | Q15643 | 228 kDa | 0 | 55 | 55 | 0 |
| Mucin-5B OS=Homo sapiens OX=9606 GN=MUC5B PE=1 SV=3 | Q9HC84 | 596 kDa | 0 | 51 | 51 | 0 |
| ARF GTPase-activating protein GIT1 OS=Homo sapiens OX=9606 GN=GIT1 PE=1 SV=2 | Q9Y2X7 | 84 kDa | 0 | 14 | 14 | 0 |
| Mucin-5AC OS=Homo sapiens OX=9606 GN=MUC5AC PE=1 SV=4 | P98088 | 586 kDa | 0 | 13 | 13 | 0 |
| BPI fold-containing family B member 1 OS=Homo sapiens OX=9606 GN=BPIFB1 PE=1 SV=1 | Q8TDL5 | 52 kDa | 0 | 12 | 12 | 0 |
| Rho guanine nucleotide exchange factor 7 OS=Homo sapiens OX=9606 GN=ARHGEF7 PE=1 SV=2 | Q14155 | 90 kDa | 0 | 10 | 10 | 0 |
| Fibronectin OS=Homo sapiens OX=9606 GN=FN1 PE=1 SV=4 | P02751 | 263 kDa | 0 | 9 | 9 | 0 |
| Haptoglobin OS=Homo sapiens OX=9606 GN=HP PE=1 SV=1 | P00738 | 45 kDa | 0 | 8 | 8 | 0 |
| BPI fold-containing family B member 4 OS=Homo sapiens OX=9606 GN=BPIFB4 PE=2 SV=2 | P59827 | 65 kDa | 0 | 8 | 8 | 0 |
| Coatomer subunit alpha OS=Homo sapiens OX=9606 GN=COPA PE=1 SV=2 | P53621 | 138 kDa | 0 | 6 | 6 | 0 |

| | | | | | | |
|---|---|---|---|---|---|---|
| Dihydrolipoyllysine-residue succinyltransferase component of 2-oxoglutarate dehydrogenase complex, mitochondrial OS=Homo sapiens OX=9606 GN=DLST PE=1 SV=4 | P36957 | 49 kDa | 1 | 12 | 11 | 1 |
| Deleted in malignant brain tumors 1 protein OS=Homo sapiens OX=9606 GN=DMBT1 PE=1 SV=2 | Q9UGM3 | 261 kDa | 1 | 11 | 10 | 1 |
| Zymogen granule protein 16 homolog B OS=Homo sapiens OX=9606 GN=ZG16B PE=1 SV=3 | Q96DA0 | 23 kDa | 0 | 5 | 5 | 0 |
| Apolipoprotein A-I OS=Homo sapiens OX=9606 GN=APOA1 PE=1 SV=1 | P02647 | 31 kDa | 0 | 5 | 5 | 0 |
| Thrombospondin-1 OS=Homo sapiens OX=9606 GN=THBS1 PE=1 SV=2 | P07996 | 129 kDa | 0 | 5 | 5 | 0 |
| Mesothelin OS=Homo sapiens OX=9606 GN=MSLN PE=1 SV=2 | Q13421 | 69 kDa | 0 | 5 | 5 | 0 |
| Annexin A1 OS=Homo sapiens OX=9606 GN=ANXA1 PE=1 SV=2 | P04083 | 39 kDa | 0 | 4 | 4 | 0 |
| Complement factor B OS=Homo sapiens OX=9606 GN=CFB PE=1 SV=2 | P00751 | 86 kDa | 0 | 4 | 4 | 0 |
| ARF GTPase-activating protein GIT2 OS=Homo sapiens OX=9606 GN=GIT2 PE=1 SV=2 | Q14161 | 85 kDa | 0 | 4 | 4 | 0 |
| Cathepsin D OS=Homo sapiens OX=9606 GN=CTSD PE=1 SV=1 | P07339 | 45 kDa | 1 | 8 | 7 | 1 |
| Serotransferrin OS=Homo sapiens OX=9606 GN=TF PE=1 SV=3 | P02787 | 77 kDa | 2 | 12 | 10 | 1 |
| Clusterin OS=Homo sapiens OX=9606 GN=CLU PE=1 SV=1 | P10909 | 52 kDa | 1 | 7 | 6 | 1 |
| Poly [ADP-ribose] polymerase 1 OS=Homo sapiens OX=9606 GN=PARP1 PE=1 SV=4 | P09874 | 113 kDa | 1 | 7 | 6 | 1 |
| Neutrophil defensin 1 OS=Homo sapiens OX=9606 GN=DEFA1 PE=1 SV=1 | P59665 | 10 kDa | 0 | 3 | 3 | 0 |
| Cornifin-B OS=Homo sapiens OX=9606 GN=SPRR1B PE=1 SV=2 | P22528 | 10 kDa | 0 | 3 | 3 | 0 |
| Ezrin OS=Homo sapiens OX=9606 GN=EZR PE=1 SV=4 | P15311 | 69 kDa | 0 | 3 | 3 | 0 |
| Cystatin-C OS=Homo sapiens OX=9606 GN=CST3 PE=1 SV=1 | P01034 | 16 kDa | 0 | 3 | 3 | 0 |
| Lipocalin-15 OS=Homo sapiens OX=9606 GN=LCN15 PE=1 SV=1 | Q6UWW0 | 20 kDa | 0 | 3 | 3 | 0 |
| Serine/threonine-protein kinase PAK 1 OS=Homo sapiens OX=9606 GN=PAK1 PE=1 SV=2 | Q13153 | 61 kDa | 0 | 3 | 3 | 0 |
| Trefoil factor 3 OS=Homo sapiens OX=9606 GN=TFF3 PE=1 SV=2 | Q07654 | 10 kDa | 0 | 3 | 3 | 0 |
| Pachytene checkpoint protein 2 homolog OS=Homo sapiens OX=9606 GN=TRIP13 PE=1 SV=2 | Q15645 | 49 kDa | 0 | 3 | 3 | 0 |
| WAP four-disulfide core domain protein 2 OS=Homo sapiens OX=9606 GN=WFDC2 PE=1 SV=2 | Q14508 | 13 kDa | 0 | 3 | 3 | 0 |

| | | | | | | |
|---|---|---|---|---|---|---|
| C-X-C motif chemokine 17 OS=Homo sapiens OX=9606 GN=CXCL17 PE=1 SV=1 | Q6UXB2 | 14 kDa | 0 | 3 | 3 | 0 |
| Protein-glutamine gamma-glutamyltransferase K OS=Homo sapiens OX=9606 GN=TGM1 PE=1 SV=4 | P22735 | 90 kDa | 0 | 3 | 3 | 0 |
| Mucin-7 OS=Homo sapiens OX=9606 GN=MUC7 PE=1 SV=2 | Q8TAX7 | 39 kDa | 0 | 3 | 3 | 0 |
| Splicing factor 3B subunit 1 OS=Homo sapiens OX=9606 GN=SF3B1 PE=1 SV=3 | O75533 | 146 kDa | 1 | 6 | 5 | 1 |
| Serpin B3 OS=Homo sapiens OX=9606 GN=SERPINB3 PE=1 SV=2 | P29508 | 45 kDa | 2 | 9 | 7 | 2 |
| Antileukoproteinase OS=Homo sapiens OX=9606 GN=SLPI PE=1 SV=2 | P03973 | 14 kDa | 2 | 8 | 6 | 2 |
| Protein S100-A9 OS=Homo sapiens OX=9606 GN=S100A9 PE=1 SV=1 | P06702 | 13 kDa | 2 | 8 | 6 | 2 |
| Junction plakoglobin OS=Homo sapiens OX=9606 GN=JUP PE=1 SV=3 | P14923 | 82 kDa | 5 | 16 | 11 | 5 |
| Complement C3 OS=Homo sapiens OX=9606 GN=C3 PE=1 SV=2 | P01024 | 187 kDa | 4 | 13 | 9 | 4 |
| Protein LEG1 homolog OS=Homo sapiens OX=9606 GN=LEG1 PE=1 SV=2 | Q6P5S2 | 38 kDa | 1 | 4 | 3 | 1 |
| 60S ribosomal protein L15 OS=Homo sapiens OX=9606 GN=RPL15 PE=1 SV=2 | P61313 | 24 kDa | 2 | 6 | 4 | 2 |
| RuvB-like 2 OS=Homo sapiens OX=9606 GN=RUVBL2 PE=1 SV=3 | Q9Y230 | 51 kDa | 2 | 6 | 4 | 2 |
| Fatty acid-binding protein, epidermal OS=Homo sapiens OX=9606 GN=FABP5 PE=1 SV=3 | Q01469 | 15 kDa | 4 | 10 | 6 | 4 |
| Desmoplakin OS=Homo sapiens OX=9606 GN=DSP PE=1 SV=3 | P15924 | 332 kDa | 17 | 38 | 21 | 17 |
| Lysozyme C OS=Homo sapiens OX=9606 GN=LYZ PE=1 SV=1 | P61626 | 17 kDa | 5 | 11 | 6 | 5 |
| Annexin A2 OS=Homo sapiens OX=9606 GN=ANXA2 PE=1 SV=2 | P07355 | 39 kDa | 3 | 7 | 4 | 3 |
| Glutaryl-CoA dehydrogenase, mitochondrial OS=Homo sapiens OX=9606 GN=GCDH PE=1 SV=1 | Q92947 | 48 kDa | 2 | 5 | 3 | 2 |
| 60S ribosomal protein L32 OS=Homo sapiens OX=9606 GN=RPL32 PE=1 SV=2 | P62910 | 16 kDa | 2 | 5 | 3 | 2 |
| Alpha-2-macroglobulin OS=Homo sapiens OX=9606 GN=A2M PE=1 SV=3 | P01023 | 163 kDa | 9 | 18 | 9 | 7 |
| 60S ribosomal protein L13a OS=Homo sapiens OX=9606 GN=RPL13A PE=1 SV=2 | P40429 | 24 kDa | 3 | 6 | 3 | 3 |
| Elongation factor Tu, mitochondrial OS=Homo sapiens OX=9606 GN=TUFM PE=1 SV=2 | P49411 | 50 kDa | 7 | 12 | 5 | 6 |

| | | | | | | |
|---|---|---|---|---|---|---|
| 60S acidic ribosomal protein P2 OS=Homo sapiens OX=9606 GN=RPLP2 PE=1 SV=1 | P05387 | 12 kDa | 4 | 7 | 3 | 3 |
| Retinitis pigmentosa 9 protein OS=Homo sapiens OX=9606 GN=RP9 PE=1 SV=2 | Q8TA86 | 26 kDa | 6 | 10 | 4 | 6 |
| Calnexin OS=Homo sapiens OX=9606 GN=CANX PE=1 SV=2 | P27824 | 68 kDa | 7 | 11 | 4 | 6 |
| Filaggrin OS=Homo sapiens OX=9606 GN=FLG PE=1 SV=3 | P20930 | 435 kDa | 10 | 15 | 5 | 10 |
| Protein LSM14 homolog A OS=Homo sapiens OX=9606 GN=LSM14A PE=1 SV=3 | Q8ND56 | 51 kDa | 7 | 10 | 3 | 6 |
| Suprabasin OS=Homo sapiens OX=9606 GN=SBSN PE=1 SV=2 | Q6UWP8 | 61 kDa | 9 | 12 | 3 | 7 |
| Lactotransferrin OS=Homo sapiens OX=9606 GN=LTF PE=1 SV=6 | P02788 | 78 kDa | 38 | 47 | 9 | 29 |

**Table S3. Proteins identified exclusively in the GST-pulldown experiment.**

| | | | Total Spectrum Count GST-Pulldown (HEK293T) | | | Exclusi GST- |
|---|---|---|---|---|---|---|
| Identified Proteins | Accession Number | Molecular Weight | CTRL | TRMT | TRMT-CTRL | CTRL |
| DNA ligase 3 OS=Homo sapiens OX=9606 GN=LIG3 PE=1 SV=2 | P49916 | 113 kDa | 0 | 33 | 33 | 0 |
| Cold shock domain-containing protein E1 OS=Homo sapiens OX=9606 GN=CSDE1 PE=1 SV=2 | O75534 | 89 kDa | 0 | 25 | 25 | 0 |
| DNA mismatch repair protein Msh6 OS=Homo sapiens OX=9606 GN=MSH6 PE=1 SV=2 | P52701 | 153 kDa | 0 | 18 | 18 | 0 |
| DNA repair protein XRCC1 OS=Homo sapiens OX=9606 GN=XRCC1 PE=1 SV=2 | P18887 | 69 kDa | 0 | 16 | 16 | 0 |
| 60S ribosomal protein L10 OS=Homo sapiens OX=9606 GN=RPL10 PE=1 SV=4 | P27635 | 25 kDa | 0 | 15 | 15 | 0 |
| ATPase family AAA domain-containing protein 3A OS=Homo sapiens OX=9606 GN=ATAD3A PE=1 SV=2 | Q9NVI7 | 71 kDa | 0 | 14 | 14 | 0 |
| Replication protein A 70 kDa DNA-binding subunit OS=Homo sapiens OX=9606 GN=RPA1 PE=1 SV=2 | P27694 | 68 kDa | 0 | 14 | 14 | 0 |
| Serine/arginine-rich splicing factor 9 OS=Homo sapiens OX=9606 GN=SRSF9 PE=1 SV=1 | Q13242 | 26 kDa | 0 | 12 | 12 | 0 |
| Probable ATP-dependent RNA helicase DDX46 OS=Homo sapiens OX=9606 GN=DDX46 PE=1 SV=2 | Q7L014 | 117 kDa | 0 | 11 | 11 | 0 |
| RNA transcription, translation and transport factor protein OS=Homo sapiens OX=9606 GN=RTRAF PE=1 SV=1 | Q9Y224 | 28 kDa | 0 | 11 | 11 | 0 |

| | | | | | | |
|---|---|---|---|---|---|---|
| DNA topoisomerase 3-alpha OS=Homo sapiens OX=9606 GN=TOP3A PE=1 SV=1 | Q13472 | 112 kDa | 0 | 11 | 11 | 0 |
| KH domain-containing, RNA-binding, signal transduction-associated protein 1 OS=Homo sapiens OX=9606 GN=KHDRBS1 PE=1 SV=1 | Q07666 | 48 kDa | 0 | 10 | 10 | 0 |
| Tripartite motif-containing protein 26 OS=Homo sapiens OX=9606 GN=TRIM26 PE=1 SV=1 | Q12899 | 62 kDa | 0 | 10 | 10 | 0 |
| Chromodomain-helicase-DNA-binding protein 1-like OS=Homo sapiens OX=9606 GN=CHD1L PE=1 SV=2 | Q86WJ1 | 101 kDa | 0 | 10 | 10 | 0 |
| Protein FAM98A OS=Homo sapiens OX=9606 GN=FAM98A PE=1 SV=1 | Q8NCA5 | 55 kDa | 0 | 10 | 10 | 0 |
| Luc7-like protein 3 OS=Homo sapiens OX=9606 GN=LUC7L3 PE=1 SV=2 | O95232 | 51 kDa | 0 | 9 | 9 | 0 |
| Structural maintenance of chromosomes protein 1A OS=Homo sapiens OX=9606 GN=SMC1A PE=1 SV=2 | Q14683 | 143 kDa | 0 | 9 | 9 | 0 |
| Protein FAM98B OS=Homo sapiens OX=9606 GN=FAM98B PE=1 SV=1 | Q52LJ0 | 37 kDa | 0 | 9 | 9 | 0 |
| Eukaryotic translation initiation factor 3 subunit D OS=Homo sapiens OX=9606 GN=EIF3D PE=1 SV=1 | O15371 | 64 kDa | 0 | 9 | 9 | 0 |
| X-ray repair cross-complementing protein 5 OS=Homo sapiens OX=9606 GN=XRCC5 PE=1 SV=3 | P13010 | 83 kDa | 0 | 9 | 9 | 0 |
| ATP-dependent RNA helicase DHX36 OS=Homo sapiens OX=9606 GN=DHX36 PE=1 SV=2 | Q9H2U1 | 115 kDa | 0 | 9 | 9 | 0 |
| Replication factor C subunit 1 OS=Homo sapiens OX=9606 GN=RFC1 PE=1 SV=4 | P35251 | 128 kDa | 0 | 9 | 9 | 0 |
| Helicase-like transcription factor OS=Homo sapiens OX=9606 GN=HLTF PE=1 SV=2 | Q14527 | 114 kDa | 0 | 9 | 9 | 0 |
| Emerin OS=Homo sapiens OX=9606 GN=EMD PE=1 SV=1 | P50402 | 29 kDa | 0 | 8 | 8 | 0 |
| Cleavage and polyadenylation specificity factor subunit 6 OS=Homo sapiens OX=9606 GN=CPSF6 PE=1 SV=2 | Q16630 | 59 kDa | 0 | 7 | 7 | 0 |
| B-cell receptor-associated protein 31 OS=Homo sapiens OX=9606 GN=BCAP31 PE=1 SV=3 | P51572 | 28 kDa | 0 | 7 | 7 | 0 |
| ATPase inhibitor, mitochondrial OS=Homo sapiens OX=9606 GN=ATPIF1 PE=1 SV=1 | Q9UII2 | 12 kDa | 0 | 7 | 7 | 0 |
| 60S ribosomal protein L26 OS=Homo sapiens OX=9606 GN=RPL26 PE=1 SV=1 | P61254 | 17 kDa | 0 | 7 | 7 | 0 |
| Bromodomain-containing protein 2 OS=Homo sapiens OX=9606 GN=BRD2 PE=1 SV=2 | P25440 | 88 kDa | 0 | 7 | 7 | 0 |

| | | | | | | |
|---|---|---|---|---|---|---|
| DNA-directed RNA polymerase III subunit RPC1 OS=Homo sapiens OX=9606 GN=POLR3A PE=1 SV=2 | O14802 | 156 kDa | 0 | 7 | 7 | 0 |
| RPA-related protein RADX OS=Homo sapiens OX=9606 GN=RADX PE=1 SV=2 | Q6NSI4 | 98 kDa | 0 | 7 | 7 | 0 |
| DNA dC->dU-editing enzyme APOBEC-3C OS=Homo sapiens OX=9606 GN=APOBEC3C PE=1 SV=2 | Q9NRW3 | 23 kDa | 0 | 7 | 7 | 0 |
| Pre-mRNA-splicing factor 38A OS=Homo sapiens OX=9606 GN=PRPF38A PE=1 SV=1 | Q8NAV1 | 37 kDa | 0 | 6 | 6 | 0 |
| Delta-1-pyrroline-5-carboxylate synthase OS=Homo sapiens OX=9606 GN=ALDH18A1 PE=1 SV=2 | P54886 | 87 kDa | 0 | 6 | 6 | 0 |
| Ribose-phosphate pyrophosphokinase 1 OS=Homo sapiens OX=9606 GN=PRPS1 PE=1 SV=2 | P60891 | 35 kDa | 0 | 6 | 6 | 0 |
| DNA polymerase beta OS=Homo sapiens OX=9606 GN=POLB PE=1 SV=3 | P06746 | 38 kDa | 0 | 6 | 6 | 0 |
| Glutaredoxin-3 OS=Homo sapiens OX=9606 GN=GLRX3 PE=1 SV=2 | O76003 | 37 kDa | 0 | 6 | 6 | 0 |
| Coiled-coil domain-containing protein 47 OS=Homo sapiens OX=9606 GN=CCDC47 PE=1 SV=1 | Q96A33 | 56 kDa | 0 | 6 | 6 | 0 |
| DNA damage-binding protein 1 OS=Homo sapiens OX=9606 GN=DDB1 PE=1 SV=1 | Q16531 | 127 kDa | 0 | 6 | 6 | 0 |
| Protein FRA10AC1 OS=Homo sapiens OX=9606 GN=FRA10AC1 PE=1 SV=3 | Q70Z53 | 38 kDa | 0 | 6 | 6 | 0 |
| Zinc finger CCCH-type antiviral protein 1 OS=Homo sapiens OX=9606 GN=ZC3HAV1 PE=1 SV=3 | Q7Z2W4 | 101 kDa | 0 | 6 | 6 | 0 |
| 5-methylcytosine rRNA methyltransferase NSUN4 OS=Homo sapiens OX=9606 GN=NSUN4 PE=1 SV=2 | Q96CB9 | 43 kDa | 0 | 6 | 6 | 0 |
| Mitochondrial genome maintenance exonuclease 1 OS=Homo sapiens OX=9606 GN=MGME1 PE=1 SV=1 | Q9BQP7 | 39 kDa | 0 | 6 | 6 | 0 |
| E3 ubiquitin-protein ligase RNF114 OS=Homo sapiens OX=9606 GN=RNF114 PE=1 SV=1 | Q9Y508 | 26 kDa | 0 | 6 | 6 | 0 |
| CWF19-like protein 2 OS=Homo sapiens OX=9606 GN=CWF19L2 PE=1 SV=4 | Q2TBE0 | 104 kDa | 0 | 5 | 5 | 0 |
| Cyclin-dependent kinase 1 OS=Homo sapiens OX=9606 GN=CDK1 PE=1 SV=3 | P06493 | 34 kDa | 0 | 5 | 5 | 0 |
| Catenin delta-1 OS=Homo sapiens OX=9606 GN=CTNND1 PE=1 SV=1 | O60716 | 108 kDa | 0 | 5 | 5 | 0 |
| Succinate--CoA ligase [ADP/GDP-forming] subunit alpha, | P53597 | 36 kDa | 0 | 5 | 5 | 0 |

| | | | | | | |
|---|---|---|---|---|---|---|
| mitochondrial OS=Homo sapiens OX=9606 GN=SUCLG1 PE=1 SV=4 | | | | | | |
| Microfibrillar-associated protein 1 OS=Homo sapiens OX=9606 GN=MFAP1 PE=1 SV=2 | P55081 | 52 kDa | 0 | 5 | 5 | 0 |
| Coiled-coil domain-containing protein 124 OS=Homo sapiens OX=9606 GN=CCDC124 PE=1 SV=1 | Q96CT7 | 26 kDa | 0 | 5 | 5 | 0 |
| Negative elongation factor E OS=Homo sapiens OX=9606 GN=NELFE PE=1 SV=3 | P18615 | 43 kDa | 0 | 5 | 5 | 0 |
| FACT complex subunit SSRP1 OS=Homo sapiens OX=9606 GN=SSRP1 PE=1 SV=1 | Q08945 | 81 kDa | 0 | 5 | 5 | 0 |
| Transcription termination factor 4, mitochondrial OS=Homo sapiens OX=9606 GN=MTERF4 PE=1 SV=3 | Q7Z6M4 | 44 kDa | 0 | 5 | 5 | 0 |
| E3 ubiquitin-protein ligase RNF138 OS=Homo sapiens OX=9606 GN=RNF138 PE=1 SV=1 | Q8WVD3 | 28 kDa | 0 | 5 | 5 | 0 |
| 5'-3' exoribonuclease 2 OS=Homo sapiens OX=9606 GN=XRN2 PE=1 SV=1 | Q9H0D6 | 109 kDa | 0 | 5 | 5 | 0 |
| Homeobox protein Hox-B6 OS=Homo sapiens OX=9606 GN=HOXB6 PE=1 SV=4 | P17509 | 25 kDa | 0 | 5 | 5 | 0 |
| DNA mismatch repair protein Msh3 OS=Homo sapiens OX=9606 GN=MSH3 PE=1 SV=4 | P20585 | 127 kDa | 0 | 5 | 5 | 0 |
| Replication protein A 14 kDa subunit OS=Homo sapiens OX=9606 GN=RPA3 PE=1 SV=1 | P35244 | 14 kDa | 0 | 5 | 5 | 0 |
| Cleavage and polyadenylation specificity factor subunit 7 OS=Homo sapiens OX=9606 GN=CPSF7 PE=1 SV=1 | Q8N684 | 52 kDa | 0 | 4 | 4 | 0 |
| Pre-mRNA-splicing factor 38B OS=Homo sapiens OX=9606 GN=PRPF38B PE=1 SV=1 | Q5VTL8 | 64 kDa | 0 | 4 | 4 | 0 |
| NF-kappa-B-activating protein OS=Homo sapiens OX=9606 GN=NKAP PE=1 SV=1 | Q8N5F7 | 47 kDa | 0 | 4 | 4 | 0 |
| Medium-chain specific acyl-CoA dehydrogenase, mitochondrial OS=Homo sapiens OX=9606 GN=ACADM PE=1 SV=1 | P11310 | 47 kDa | 0 | 4 | 4 | 0 |
| Protein FRG1 OS=Homo sapiens OX=9606 GN=FRG1 PE=1 SV=1 | Q14331 | 29 kDa | 0 | 4 | 4 | 0 |
| Serine/arginine-rich splicing factor 4 OS=Homo sapiens OX=9606 GN=SRSF4 PE=1 SV=2 | Q08170 | 57 kDa | 0 | 4 | 4 | 0 |
| Deoxynucleoside triphosphate triphosphohydrolase SAMHD1 OS=Homo sapiens OX=9606 GN=SAMHD1 PE=1 SV=2 | Q9Y3Z3 | 72 kDa | 0 | 4 | 4 | 0 |
| DNA mismatch repair protein Msh2 OS=Homo sapiens OX=9606 GN=MSH2 PE=1 SV=1 | P43246 | 105 kDa | 0 | 4 | 4 | 0 |

| | | | | | | |
|---|---|---|---|---|---|---|
| rRNA 2'-O-methyltransferase fibrillarin OS=Homo sapiens OX=9606 GN=FBL PE=1 SV=2 | P22087 | 34 kDa | 0 | 4 | 4 | 0 |
| Protein FAM32A OS=Homo sapiens OX=9606 GN=FAM32A PE=1 SV=2 | Q9Y421 | 13 kDa | 0 | 4 | 4 | 0 |
| 60S ribosomal protein L35 OS=Homo sapiens OX=9606 GN=RPL35 PE=1 SV=2 | P42766 | 15 kDa | 0 | 4 | 4 | 0 |
| Mitochondrial import inner membrane translocase subunit TIM50 OS=Homo sapiens OX=9606 GN=TIMM50 PE=1 SV=2 | Q3ZCQ8 | 40 kDa | 0 | 4 | 4 | 0 |
| Multiple myeloma tumor-associated protein 2 OS=Homo sapiens OX=9606 GN=MMTAG2 PE=1 SV=1 | Q9BU76 | 29 kDa | 0 | 4 | 4 | 0 |
| Calcium-binding mitochondrial carrier protein Aralar2 OS=Homo sapiens OX=9606 GN=SLC25A13 PE=1 SV=2 | Q9UJS0 | 74 kDa | 0 | 4 | 4 | 0 |
| Elongin-A OS=Homo sapiens OX=9606 GN=ELOA PE=1 SV=2 | Q14241 | 90 kDa | 0 | 4 | 4 | 0 |
| DnaJ homolog subfamily C member 7 OS=Homo sapiens OX=9606 GN=DNAJC7 PE=1 SV=2 | Q99615 | 56 kDa | 0 | 4 | 4 | 0 |
| Protein pelota homolog OS=Homo sapiens OX=9606 GN=PELO PE=1 SV=2 | Q9BRX2 | 43 kDa | 0 | 4 | 4 | 0 |
| YTH domain-containing family protein 2 OS=Homo sapiens OX=9606 GN=YTHDF2 PE=1 SV=2 | Q9Y5A9 | 62 kDa | 0 | 4 | 4 | 0 |
| Translation machinery-associated protein 16 OS=Homo sapiens OX=9606 GN=TMA16 PE=1 SV=2 | Q96EY4 | 24 kDa | 0 | 4 | 4 | 0 |
| Major centromere autoantigen B OS=Homo sapiens OX=9606 GN=CENPB PE=1 SV=2 | P07199 | 65 kDa | 0 | 4 | 4 | 0 |
| DNA polymerase delta catalytic subunit OS=Homo sapiens OX=9606 GN=POLD1 PE=1 SV=2 | P28340 | 124 kDa | 0 | 4 | 4 | 0 |
| DNA-directed RNA polymerase II subunit RPB2 OS=Homo sapiens OX=9606 GN=POLR2B PE=1 SV=1 | P30876 | 134 kDa | 0 | 4 | 4 | 0 |
| Homeobox protein Hox-A11 OS=Homo sapiens OX=9606 GN=HOXA11 PE=2 SV=2 | P31270 | 34 kDa | 0 | 4 | 4 | 0 |
| Cleavage stimulation factor subunit 1 OS=Homo sapiens OX=9606 GN=CSTF1 PE=1 SV=1 | Q05048 | 48 kDa | 0 | 4 | 4 | 0 |
| RING finger protein 166 OS=Homo sapiens OX=9606 GN=RNF166 PE=2 SV=1 | Q96A37 | 26 kDa | 0 | 4 | 4 | 0 |
| RNA-binding protein MEX3A OS=Homo sapiens OX=9606 GN=MEX3A PE=1 SV=1 | A1L020 | 54 kDa | 0 | 4 | 4 | 0 |
| Replication protein A 32 kDa subunit OS=Homo sapiens OX=9606 GN=RPA2 PE=1 SV=1 | P15927 | 29 kDa | 0 | 4 | 4 | 0 |

| | | | | | | |
|---|---|---|---|---|---|---|
| Homeobox protein Hox-C9 OS=Homo sapiens OX=9606 GN=HOXC9 PE=1 SV=3 | P31274 | 29 kDa | 0 | 4 | 4 | 0 |
| Polyglutamine-binding protein 1 OS=Homo sapiens OX=9606 GN=PQBP1 PE=1 SV=1 | O60828 | 30 kDa | 0 | 4 | 4 | 0 |
| DDB1- and CUL4-associated factor 7 OS=Homo sapiens OX=9606 GN=DCAF7 PE=1 SV=1 | P61962 | 39 kDa | 0 | 4 | 4 | 0 |
| Lysozyme C OS=Homo sapiens OX=9606 GN=LYZ PE=1 SV=1 | P61626 | 17 kDa | 0 | 4 | 4 | 0 |
| Pre-mRNA-splicing factor CWC22 homolog OS=Homo sapiens OX=9606 GN=CWC22 PE=1 SV=3 | Q9HCG8 | 105 kDa | 0 | 3 | 3 | 0 |
| U2 snRNP-associated SURP motif-containing protein OS=Homo sapiens OX=9606 GN=U2SURP PE=1 SV=2 | O15042 | 118 kDa | 0 | 3 | 3 | 0 |
| 60S ribosomal protein L37a OS=Homo sapiens OX=9606 GN=RPL37A PE=1 SV=2 | P61513 | 10 kDa | 0 | 3 | 3 | 0 |
| Serine/Arginine-related protein 53 OS=Homo sapiens OX=9606 GN=RSRC1 PE=1 SV=1 | Q96IZ7 | 39 kDa | 0 | 3 | 3 | 0 |
| RNA-binding protein 39 OS=Homo sapiens OX=9606 GN=RBM39 PE=1 SV=2 | Q14498 | 59 kDa | 0 | 3 | 3 | 0 |
| Pre-mRNA-splicing factor CWC25 homolog OS=Homo sapiens OX=9606 GN=CWC25 PE=1 SV=1 | Q9NXE8 | 50 kDa | 0 | 3 | 3 | 0 |
| 40S ribosomal protein S23 OS=Homo sapiens OX=9606 GN=RPS23 PE=1 SV=3 | P62266 | 16 kDa | 0 | 3 | 3 | 0 |
| Translation machinery-associated protein 7 OS=Homo sapiens OX=9606 GN=TMA7 PE=1 SV=1 | Q9Y2S6 | 7 kDa | 0 | 3 | 3 | 0 |
| Serine/arginine-rich splicing factor 11 OS=Homo sapiens OX=9606 GN=SRSF11 PE=1 SV=1 | Q05519 | 54 kDa | 0 | 3 | 3 | 0 |
| DNA-dependent protein kinase catalytic subunit OS=Homo sapiens OX=9606 GN=PRKDC PE=1 SV=3 | P78527 | 469 kDa | 0 | 3 | 3 | 0 |
| Trifunctional enzyme subunit beta, mitochondrial OS=Homo sapiens OX=9606 GN=HADHB PE=1 SV=3 | P55084 | 51 kDa | 0 | 3 | 3 | 0 |
| Cytochrome c oxidase subunit NDUFA4 OS=Homo sapiens OX=9606 GN=NDUFA4 PE=1 SV=1 | O00483 | 9 kDa | 0 | 3 | 3 | 0 |
| 28S ribosomal protein S12, mitochondrial OS=Homo sapiens OX=9606 GN=MRPS12 PE=1 SV=1 | O15235 | 15 kDa | 0 | 3 | 3 | 0 |
| Protein DEK OS=Homo sapiens OX=9606 GN=DEK PE=1 SV=1 | P35659 | 43 kDa | 0 | 3 | 3 | 0 |
| Non-histone chromosomal protein HMG-17 OS=Homo sapiens OX=9606 GN=HMGN2 PE=1 SV=3 | P05204 | 9 kDa | 0 | 3 | 3 | 0 |
| NADH dehydrogenase [ubiquinone] 1 alpha subcomplex subunit 10, | O95299 | 41 kDa | 0 | 3 | 3 | 0 |

| | | | | | | |
|---|---|---|---|---|---|---|
| mitochondrial OS=Homo sapiens OX=9606 GN=NDUFA10 PE=1 SV=1 | | | | | | |
| Replication factor C subunit 3 OS=Homo sapiens OX=9606 GN=RFC3 PE=1 SV=2 | P40938 | 41 kDa | 0 | 3 | 3 | 0 |
| 40S ribosomal protein S15 OS=Homo sapiens OX=9606 GN=RPS15 PE=1 SV=2 | P62841 | 17 kDa | 0 | 3 | 3 | 0 |
| Trifunctional enzyme subunit alpha, mitochondrial OS=Homo sapiens OX=9606 GN=HADHA PE=1 SV=2 | P40939 | 83 kDa | 0 | 3 | 3 | 0 |
| Corepressor interacting with RBPJ 1 OS=Homo sapiens OX=9606 GN=CIR1 PE=1 SV=1 | Q86X95 | 52 kDa | 0 | 3 | 3 | 0 |
| Transducin beta-like protein 2 OS=Homo sapiens OX=9606 GN=TBL2 PE=1 SV=1 | Q9Y4P3 | 50 kDa | 0 | 3 | 3 | 0 |
| Zinc finger matrin-type protein 2 OS=Homo sapiens OX=9606 GN=ZMAT2 PE=1 SV=1 | Q96NC0 | 24 kDa | 0 | 3 | 3 | 0 |
| HBS1-like protein OS=Homo sapiens OX=9606 GN=HBS1L PE=1 SV=1 | Q9Y450 | 75 kDa | 0 | 3 | 3 | 0 |
| RNA/RNP complex-1-interacting phosphatase OS=Homo sapiens OX=9606 GN=DUSP11 PE=1 SV=2 | O75319 | 44 kDa | 0 | 3 | 3 | 0 |
| tRNA methyltransferase 10 homolog C OS=Homo sapiens OX=9606 GN=TRMT10C PE=1 SV=2 | Q7L0Y3 | 47 kDa | 0 | 3 | 3 | 0 |
| DNA-directed RNA polymerase III subunit RPC4 OS=Homo sapiens OX=9606 GN=POLR3D PE=1 SV=2 | P05423 | 44 kDa | 0 | 3 | 3 | 0 |
| GTPase Era, mitochondrial OS=Homo sapiens OX=9606 GN=ERAL1 PE=1 SV=2 | O75616 | 48 kDa | 0 | 3 | 3 | 0 |
| Homeobox protein Hox-B8 OS=Homo sapiens OX=9606 GN=HOXB8 PE=2 SV=2 | P17481 | 28 kDa | 0 | 3 | 3 | 0 |
| Replication factor C subunit 4 OS=Homo sapiens OX=9606 GN=RFC4 PE=1 SV=2 | P35249 | 40 kDa | 0 | 3 | 3 | 0 |
| DNA-directed RNA polymerases I, II, and III subunit RPABC3 OS=Homo sapiens OX=9606 GN=POLR2H PE=1 SV=4 | P52434 | 17 kDa | 0 | 3 | 3 | 0 |
| N-alpha-acetyltransferase 40 OS=Homo sapiens OX=9606 GN=NAA40 PE=1 SV=1 | Q86UY6 | 27 kDa | 0 | 3 | 3 | 0 |
| Histone H1x OS=Homo sapiens OX=9606 GN=H1FX PE=1 SV=1 | Q92522 | 22 kDa | 0 | 3 | 3 | 0 |
| HIV Tat-specific factor 1 OS=Homo sapiens OX=9606 GN=HTATSF1 PE=1 SV=1 | O43719 | 86 kDa | 0 | 3 | 3 | 0 |
| Forkhead box protein C1 OS=Homo sapiens OX=9606 GN=FOXC1 PE=1 SV=3 | Q12948 | 57 kDa | 0 | 3 | 3 | 0 |
| Homeobox protein Hox-A5 OS=Homo sapiens OX=9606 GN=HOXA5 PE=1 SV=2 | P20719 | 29 kDa | 0 | 3 | 3 | 0 |

| | | | | | | |
|---|---|---|---|---|---|---|
| DNA-directed RNA polymerase III subunit RPC2 OS=Homo sapiens OX=9606 GN=POLR3B PE=1 SV=2 | Q9NW08 | 128 kDa | 0 | 3 | 3 | 0 |
| Nucleolar protein 56 OS=Homo sapiens OX=9606 GN=NOP56 PE=1 SV=4 | O00567 | 66 kDa | 0 | 3 | 3 | 0 |
| DDB1- and CUL4-associated factor 11 OS=Homo sapiens OX=9606 GN=DCAF11 PE=1 SV=1 | Q8TEB1 | 62 kDa | 0 | 3 | 3 | 0 |
| RNA-binding protein 3 OS=Homo sapiens OX=9606 GN=RBM3 PE=1 SV=1 | P98179 | 17 kDa | 0 | 3 | 3 | 0 |
| E3 ubiquitin-protein ligase MIB2 OS=Homo sapiens OX=9606 GN=MIB2 PE=1 SV=3 | Q96AX9 | 110 kDa | 0 | 3 | 3 | 0 |
| DNA-directed RNA polymerases I and III subunit RPAC1 OS=Homo sapiens OX=9606 GN=POLR1C PE=1 SV=1 | O15160 | 39 kDa | 0 | 3 | 3 | 0 |
| Retinitis pigmentosa 9 protein OS=Homo sapiens OX=9606 GN=RP9 PE=1 SV=2 | Q8TA86 | 26 kDa | 0 | 3 | 3 | 0 |
| Poly [ADP-ribose] polymerase 2 OS=Homo sapiens OX=9606 GN=PARP2 PE=1 SV=2 | Q9UGN5 | 66 kDa | 1 | 35 | 34 | 1 |
| ATP-dependent RNA helicase DDX1 OS=Homo sapiens OX=9606 GN=DDX1 PE=1 SV=2 | Q92499 | 82 kDa | 1 | 26 | 25 | 1 |
| Activated RNA polymerase II transcriptional coactivator p15 OS=Homo sapiens OX=9606 GN=SUB1 PE=1 SV=3 | P53999 | 14 kDa | 1 | 22 | 21 | 1 |
| ATP-dependent RNA helicase DDX3X OS=Homo sapiens OX=9606 GN=DDX3X PE=1 SV=3 | O00571 | 73 kDa | 4 | 49 | 45 | 4 |
| Insulin receptor substrate 4 OS=Homo sapiens OX=9606 GN=IRS4 PE=1 SV=1 | O14654 | 134 kDa | 1 | 18 | 17 | 1 |
| tRNA-splicing ligase RtcB homolog OS=Homo sapiens OX=9606 GN=RTCB PE=1 SV=1 | Q9Y3I0 | 55 kDa | 2 | 26 | 24 | 2 |
| Heterogeneous nuclear ribonucleoprotein U-like protein 1 OS=Homo sapiens OX=9606 GN=HNRNPUL1 PE=1 SV=2 | Q9BUJ2 | 96 kDa | 1 | 16 | 15 | 1 |
| ATP-dependent DNA helicase Q1 OS=Homo sapiens OX=9606 GN=RECQL PE=1 SV=3 | P46063 | 73 kDa | 1 | 16 | 15 | 1 |
| Poly [ADP-ribose] polymerase 1 OS=Homo sapiens OX=9606 GN=PARP1 PE=1 SV=4 | P09874 | 113 kDa | 18 | 158 | 140 | 14 |
| Filaggrin OS=Homo sapiens OX=9606 GN=FLG PE=1 SV=3 | P20930 | 435 kDa | 1 | 14 | 13 | 1 |
| Probable ATP-dependent RNA helicase DDX5 OS=Homo sapiens OX=9606 GN=DDX5 PE=1 SV=1 | P17844 | 69 kDa | 3 | 27 | 24 | 1 |
| Protein RCC2 OS=Homo sapiens OX=9606 GN=RCC2 PE=1 SV=2 | Q9P258 | 56 kDa | 1 | 13 | 12 | 1 |

| | | | | | | |
|---|---|---|---|---|---|---|
| Nuclear ubiquitous casein and cyclin-dependent kinase substrate 1 OS=Homo sapiens OX=9606 GN=NUCKS1 PE=1 SV=1 | Q9H1E3 | 27 kDa | 3 | 26 | 23 | 3 |
| Peptidyl-prolyl cis-trans isomerase-like 4 OS=Homo sapiens OX=9606 GN=PPIL4 PE=1 SV=1 | Q8WUA2 | 57 kDa | 4 | 32 | 28 | 4 |
| Splicing factor U2AF 35 kDa subunit-like protein OS=Homo sapiens OX=9606 GN=U2AF1L5 PE=1 SV=1 | P0DN76 | 28 kDa | 1 | 12 | 11 | 1 |
| X-ray repair cross-complementing protein 6 OS=Homo sapiens OX=9606 GN=XRCC6 PE=1 SV=2 | P12956 | 70 kDa | 1 | 12 | 11 | 1 |
| 60S ribosomal protein L4 OS=Homo sapiens OX=9606 GN=RPL4 PE=1 SV=5 | P36578 | 48 kDa | 2 | 16 | 14 | 2 |
| ATP-dependent RNA helicase A OS=Homo sapiens OX=9606 GN=DHX9 PE=1 SV=4 | Q08211 | 141 kDa | 3 | 21 | 18 | 3 |
| 60S ribosomal protein L10a OS=Homo sapiens OX=9606 GN=RPL10A PE=1 SV=2 | P62906 | 25 kDa | 1 | 10 | 9 | 1 |
| Hornerin OS=Homo sapiens OX=9606 GN=HRNR PE=1 SV=2 | Q86YZ3 | 282 kDa | 6 | 37 | 31 | 5 |
| 60S ribosomal protein L3 OS=Homo sapiens OX=9606 GN=RPL3 PE=1 SV=2 | P39023 | 46 kDa | 1 | 9 | 8 | 1 |
| Heterogeneous nuclear ribonucleoprotein K OS=Homo sapiens OX=9606 GN=HNRNPK PE=1 SV=1 | P61978 | 51 kDa | 6 | 33 | 27 | 5 |
| 60S ribosomal protein L6 OS=Homo sapiens OX=9606 GN=RPL6 PE=1 SV=3 | Q02878 | 33 kDa | 2 | 13 | 11 | 2 |
| DnaJ homolog subfamily A member 1 OS=Homo sapiens OX=9606 GN=DNAJA1 PE=1 SV=2 | P31689 | 45 kDa | 3 | 17 | 14 | 3 |
| Splicing factor U2AF 65 kDa subunit OS=Homo sapiens OX=9606 GN=U2AF2 PE=1 SV=4 | P26368 | 54 kDa | 1 | 8 | 7 | 1 |
| DnaJ homolog subfamily A member 2 OS=Homo sapiens OX=9606 GN=DNAJA2 PE=1 SV=1 | O60884 | 46 kDa | 1 | 8 | 7 | 1 |
| C-1-tetrahydrofolate synthase, cytoplasmic OS=Homo sapiens OX=9606 GN=MTHFD1 PE=1 SV=3 | P11586 | 102 kDa | 4 | 21 | 17 | 4 |
| ADP/ATP translocase 2 OS=Homo sapiens OX=9606 GN=SLC25A5 PE=1 SV=7 | P05141 | 33 kDa | 3 | 16 | 13 | 3 |
| ADP-ribosylation factor-like protein 6-interacting protein 4 OS=Homo sapiens OX=9606 GN=ARL6IP4 PE=1 SV=2 | Q66PJ3 | 45 kDa | 3 | 16 | 13 | 3 |
| Arginine/serine-rich coiled-coil protein 2 OS=Homo sapiens OX=9606 GN=RSRC2 PE=1 SV=1 | Q7L4I2 | 51 kDa | 1 | 7 | 6 | 1 |

| | | | | | | |
|---|---|---|---|---|---|---|
| Lamina-associated polypeptide 2, isoforms beta/gamma OS=Homo sapiens OX=9606 GN=TMPO PE=1 SV=2 | P42167 | 51 kDa | 1 | 7 | 6 | 1 |
| Elongation factor 1-beta OS=Homo sapiens OX=9606 GN=EEF1B2 PE=1 SV=3 | P24534 | 25 kDa | 1 | 7 | 6 | 1 |
| Probable ATP-dependent RNA helicase DDX17 OS=Homo sapiens OX=9606 GN=DDX17 PE=1 SV=2 | Q92841 | 80 kDa | 7 | 29 | 22 | 6 |
| 60S ribosomal protein L38 OS=Homo sapiens OX=9606 GN=RPL38 PE=1 SV=2 | P63173 | 8 kDa | 3 | 14 | 11 | 2 |
| Suprabasin OS=Homo sapiens OX=9606 GN=SBSN PE=1 SV=2 | Q6UWP8 | 61 kDa | 2 | 10 | 8 | 2 |
| Peptidyl-prolyl cis-trans isomerase B OS=Homo sapiens OX=9606 GN=PPIB PE=1 SV=2 | P23284 | 24 kDa | 3 | 13 | 10 | 3 |
| Poly(rC)-binding protein 1 OS=Homo sapiens OX=9606 GN=PCBP1 PE=1 SV=2 | Q15365 | 37 kDa | 1 | 6 | 5 | 1 |
| 40S ribosomal protein S27 OS=Homo sapiens OX=9606 GN=RPS27 PE=1 SV=3 | P42677 | 9 kDa | 1 | 6 | 5 | 1 |
| Dolichol-phosphate mannosyltransferase subunit 1 OS=Homo sapiens OX=9606 GN=DPM1 PE=1 SV=1 | O60762 | 30 kDa | 1 | 6 | 5 | 1 |
| 60S ribosomal protein L36a OS=Homo sapiens OX=9606 GN=RPL36A PE=1 SV=2 | P83881 | 12 kDa | 1 | 6 | 5 | 1 |
| 60S ribosomal protein L13 OS=Homo sapiens OX=9606 GN=RPL13 PE=1 SV=4 | P26373 | 24 kDa | 2 | 9 | 7 | 1 |
| 60S ribosomal protein L23a OS=Homo sapiens OX=9606 GN=RPL23A PE=1 SV=1 | P62750 | 18 kDa | 2 | 9 | 7 | 2 |
| RuvB-like 2 OS=Homo sapiens OX=9606 GN=RUVBL2 PE=1 SV=3 | Q9Y230 | 51 kDa | 2 | 9 | 7 | 2 |
| Elongation factor 1-gamma OS=Homo sapiens OX=9606 GN=EEF1G PE=1 SV=3 | P26641 | 50 kDa | 11 | 38 | 27 | 9 |
| Heterogeneous nuclear ribonucleoprotein F OS=Homo sapiens OX=9606 GN=HNRNPF PE=1 SV=3 | P52597 | 46 kDa | 15 | 47 | 32 | 6 |
| Heterogeneous nuclear ribonucleoprotein U OS=Homo sapiens OX=9606 GN=HNRNPU PE=1 SV=6 | Q00839 | 91 kDa | 3 | 11 | 8 | 2 |
| 40S ribosomal protein S3a OS=Homo sapiens OX=9606 GN=RPS3A PE=1 SV=2 | P61247 | 30 kDa | 2 | 8 | 6 | 2 |
| DNA topoisomerase 1 OS=Homo sapiens OX=9606 GN=TOP1 PE=1 SV=2 | P11387 | 91 kDa | 2 | 8 | 6 | 1 |

| | | | | | | |
|---|---|---|---|---|---|---|
| 40S ribosomal protein S15a OS=Homo sapiens OX=9606 GN=RPS15A PE=1 SV=2 | P62244 | 15 kDa | 2 | 8 | 6 | 2 |
| Cytochrome b-c1 complex subunit 2, mitochondrial OS=Homo sapiens OX=9606 GN=UQCRC2 PE=1 SV=3 | P22695 | 48 kDa | 1 | 5 | 4 | 1 |
| Midkine OS=Homo sapiens OX=9606 GN=MDK PE=1 SV=1 | P21741 | 16 kDa | 1 | 5 | 4 | 1 |
| 60S acidic ribosomal protein P0 OS=Homo sapiens OX=9606 GN=RPLP0 PE=1 SV=1 | P05388 | 34 kDa | 1 | 5 | 4 | 1 |
| Acylglycerol kinase, mitochondrial OS=Homo sapiens OX=9606 GN=AGK PE=1 SV=2 | Q53H12 | 47 kDa | 1 | 5 | 4 | 1 |
| PHD finger protein 6 OS=Homo sapiens OX=9606 GN=PHF6 PE=1 SV=1 | Q8IWS0 | 41 kDa | 1 | 5 | 4 | 1 |
| Carbonyl reductase [NADPH] 1 OS=Homo sapiens OX=9606 GN=CBR1 PE=1 SV=3 | P16152 | 30 kDa | 17 | 49 | 32 | 10 |
| Zinc finger Ran-binding domain-containing protein 2 OS=Homo sapiens OX=9606 GN=ZRANB2 PE=1 SV=2 | O95218 | 37 kDa | 7 | 21 | 14 | 7 |
| Serine/arginine-rich splicing factor 2 OS=Homo sapiens OX=9606 GN=SRSF2 PE=1 SV=4 | Q01130 | 25 kDa | 3 | 10 | 7 | 2 |
| Putative RNA-binding protein Luc7-like 2 OS=Homo sapiens OX=9606 GN=LUC7L2 PE=1 SV=2 | Q9Y383 | 47 kDa | 10 | 29 | 19 | 5 |
| 60S ribosomal protein L17 OS=Homo sapiens OX=9606 GN=RPL17 PE=1 SV=3 | P18621 | 21 kDa | 2 | 7 | 5 | 2 |
| 60S ribosomal protein L11 OS=Homo sapiens OX=9606 GN=RPL11 PE=1 SV=2 | P62913 | 20 kDa | 2 | 7 | 5 | 2 |
| 60S ribosomal protein L28 OS=Homo sapiens OX=9606 GN=RPL28 PE=1 SV=3 | P46779 | 16 kDa | 2 | 7 | 5 | 2 |
| 60S ribosomal protein L18 OS=Homo sapiens OX=9606 GN=RPL18 PE=1 SV=2 | Q07020 | 22 kDa | 2 | 7 | 5 | 2 |
| 60S ribosomal protein L27a OS=Homo sapiens OX=9606 GN=RPL27A PE=1 SV=2 | P46776 | 17 kDa | 2 | 7 | 5 | 2 |
| Putative RNA-binding protein Luc7-like 1 OS=Homo sapiens OX=9606 GN=LUC7L PE=1 SV=1 | Q9NQ29 | 44 kDa | 4 | 12 | 8 | 1 |
| Serine-threonine kinase receptor-associated protein OS=Homo sapiens OX=9606 GN=STRAP PE=1 SV=1 | Q9Y3F4 | 38 kDa | 3 | 9 | 6 | 3 |
| Nucleolar and coiled-body phosphoprotein 1 OS=Homo sapiens OX=9606 GN=NOLC1 PE=1 SV=2 | Q14978 | 74 kDa | 1 | 4 | 3 | 1 |
| Serine/arginine repetitive matrix protein 2 OS=Homo sapiens OX=9606 GN=SRRM2 PE=1 SV=2 | Q9UQ35 | 300 kDa | 1 | 4 | 3 | 1 |

| | | | | | | |
|---|---|---|---|---|---|---|
| GTP-binding protein 2 OS=Homo sapiens OX=9606 GN=GTPBP2 PE=1 SV=1 | Q9BX10 | 66 kDa | 1 | 4 | 3 | 1 |
| ELAV-like protein 1 OS=Homo sapiens OX=9606 GN=ELAVL1 PE=1 SV=2 | Q15717 | 36 kDa | 1 | 4 | 3 | 1 |
| Double-stranded RNA-binding protein Staufen homolog 1 OS=Homo sapiens OX=9606 GN=STAU1 PE=1 SV=2 | O95793 | 63 kDa | 1 | 4 | 3 | 1 |
| High mobility group protein B1 OS=Homo sapiens OX=9606 GN=HMGB1 PE=1 SV=3 | P09429 | 25 kDa | 2 | 6 | 4 | 2 |
| Poly(rC)-binding protein 2 OS=Homo sapiens OX=9606 GN=PCBP2 PE=1 SV=1 | Q15366 | 39 kDa | 2 | 6 | 4 | 1 |
| 40S ribosomal protein S2 OS=Homo sapiens OX=9606 GN=RPS2 PE=1 SV=2 | P15880 | 31 kDa | 3 | 8 | 5 | 3 |
| RNA-binding motif protein, X chromosome OS=Homo sapiens OX=9606 GN=RBMX PE=1 SV=3 | P38159 | 42 kDa | 3 | 8 | 5 | 3 |
| Eukaryotic translation initiation factor 1A, X-chromosomal OS=Homo sapiens OX=9606 GN=EIF1AX PE=1 SV=2 | P47813 | 16 kDa | 14 | 30 | 16 | 5 |
| Heterogeneous nuclear ribonucleoprotein H OS=Homo sapiens OX=9606 GN=HNRNPH1 PE=1 SV=4 | P31943 | 49 kDa | 13 | 27 | 14 | 8 |
| U4/U6.U5 small nuclear ribonucleoprotein 27 kDa protein OS=Homo sapiens OX=9606 GN=SNRNP27 PE=1 SV=1 | Q8WVK2 | 19 kDa | 7 | 15 | 8 | 5 |
| Serine/arginine-rich splicing factor 3 OS=Homo sapiens OX=9606 GN=SRSF3 PE=1 SV=1 | P84103 | 19 kDa | 4 | 9 | 5 | 4 |
| 40S ribosomal protein S4, X isoform OS=Homo sapiens OX=9606 GN=RPS4X PE=1 SV=2 | P62701 | 30 kDa | 4 | 9 | 5 | 4 |
| 60S ribosomal protein L22 OS=Homo sapiens OX=9606 GN=RPL22 PE=1 SV=2 | P35268 | 15 kDa | 4 | 9 | 5 | 3 |
| 40S ribosomal protein S11 OS=Homo sapiens OX=9606 GN=RPS11 PE=1 SV=3 | P62280 | 18 kDa | 3 | 7 | 4 | 2 |
| 40S ribosomal protein S16 OS=Homo sapiens OX=9606 GN=RPS16 PE=1 SV=2 | P62249 | 16 kDa | 2 | 5 | 3 | 2 |
| Serine/arginine-rich splicing factor 6 OS=Homo sapiens OX=9606 GN=SRSF6 PE=1 SV=2 | Q13247 | 40 kDa | 2 | 5 | 3 | 2 |
| Heterogeneous nuclear ribonucleoprotein M OS=Homo sapiens OX=9606 GN=HNRNPM PE=1 SV=3 | P52272 | 78 kDa | 2 | 5 | 3 | 2 |

| | | | | | | |
|---|---|---|---|---|---|---|
| 60S ribosomal protein L24 OS=Homo sapiens OX=9606 GN=RPL24 PE=1 SV=1 | P83731 | 18 kDa | 2 | 5 | 3 | 2 |
| 60S ribosomal protein L32 OS=Homo sapiens OX=9606 GN=RPL32 PE=1 SV=2 | P62910 | 16 kDa | 2 | 5 | 3 | 2 |
| 60S ribosomal protein L15 OS=Homo sapiens OX=9606 GN=RPL15 PE=1 SV=2 | P61313 | 24 kDa | 5 | 10 | 5 | 4 |
| Desmoplakin OS=Homo sapiens OX=9606 GN=DSP PE=1 SV=3 | P15924 | 332 kDa | 22 | 41 | 19 | 20 |
| Eukaryotic translation initiation factor 5A-1 OS=Homo sapiens OX=9606 GN=EIF5A PE=1 SV=2 | P63241 | 17 kDa | 4 | 8 | 4 | 4 |
| Cellular nucleic acid-binding protein OS=Homo sapiens OX=9606 GN=CNBP PE=1 SV=1 | P62633 | 19 kDa | 4 | 8 | 4 | 4 |
| 60S ribosomal protein L23 OS=Homo sapiens OX=9606 GN=RPL23 PE=1 SV=1 | P62829 | 15 kDa | 7 | 12 | 5 | 4 |
| 40S ribosomal protein S8 OS=Homo sapiens OX=9606 GN=RPS8 PE=1 SV=2 | P62241 | 24 kDa | 4 | 7 | 3 | 4 |
| Junction plakoglobin OS=Homo sapiens OX=9606 GN=JUP PE=1 SV=3 | P14923 | 82 kDa | 11 | 18 | 7 | 11 |
| ATP synthase subunit alpha, mitochondrial OS=Homo sapiens OX=9606 GN=ATP5F1A PE=1 SV=1 | P25705 | 60 kDa | 8 | 12 | 4 | 7 |
| Eukaryotic initiation factor 4A-I OS=Homo sapiens OX=9606 GN=EIF4A1 PE=1 SV=1 | P60842 | 46 kDa | 8 | 12 | 4 | 6 |
| Histone H1.2 OS=Homo sapiens OX=9606 GN=HIST1H1C PE=1 SV=2 | P16403 | 21 kDa | 6 | 9 | 3 | 5 |
